# Varying recombination landscapes between individuals are driven by polymorphic transposable elements

**DOI:** 10.1101/2024.09.17.613564

**Authors:** Yuheng Huang, Yi Gao, Kayla Ly, Leila Lin, Jan Paul Lambooij, Elizabeth G. King, Aniek Janssen, Kevin H.-C. Wei, Yuh Chwen G. Lee

**Author notes:** Author for Correspondence: Grace Yuh Chwen Lee, Department of Ecology and Evolutionary Biology Center for Complex Biological Systems University of California, Irvine.

## Abstract

Meiotic recombination is a prominent force shaping genome evolution, and understanding the causes for varying recombination landscapes within and between species has remained a central, though challenging, question. Recombination rates are widely observed to negatively associate with the abundance of transposable elements (TEs), selfish genetic elements that move between genomic locations. While such associations are usually interpreted as recombination influencing the efficacy of selection at removing TEs, accumulating findings suggest that TEs could instead be the cause rather than the consequence. To test this prediction, we formally investigated the influence of polymorphic, putatively active TEs on recombination rates. We developed and benchmarked a novel approach that uses PacBio long-read sequencing to efficiently, accurately, and cost-effectively identify crossovers (COs), a key recombination product, among large numbers of pooled recombinant individuals. By applying this approach to Drosophila strains with distinct TE insertion profiles, we found that polymorphic TEs, especially RNA-based TEs and TEs with local enrichment of repressive marks, reduce the occurrence of COs. Such an effect leads to different CO frequencies between homologous sequences with and without TEs, contributing to varying CO maps between individuals. The suppressive effect of TEs on CO is further supported by two orthogonal approaches–analyzing the distributions of COs in panels of recombinant inbred lines in relation to TE polymorphism and applying marker-assisted estimations of CO frequencies to isogenic strains with and without transgenically inserted TEs. Our investigations reveal how the constantly changing mobilome can actively modify recombination landscapes, shaping genome evolution within and between species.

## Introduction

Homologous recombination, the exchange of genetic information between homologous chromosomes in sexually reproducing organisms, is a prominent force shaping genome evolution. During meiosis, recombination is initiated by double-strand breaks (DSBs). When DSBs are repaired using homologous chromosomes and resolved into crossovers (COs), large blocks of homologous chromosomes are reciprocally exchanged, shuffling neighboring alleles into new combinations (Zelkowski et al. 2019). This process allows for the purging of deleterious mutations (Muller 1964; Kondrashov 1988), facilitates adaptive evolution by bringing together beneficial variants (Fisher 1930; Muller 1932), and influences the efficacy of natural selection (Hill and Robertson 1966; Felsenstein 1974). Given the critical evolutionary role of recombination, estimation of its rate and distribution within genomes has long been an important research focus, as far back as (Sturtevant 1913). With decades of work, a clear pattern has emerged: both the rates of recombination and the distribution of recombination events within a genome vary significantly across taxa (Stapley et al. 2017), species (Smukowski and Noor 2011), populations (Samuk et al. 2020), and even among individuals of the same species (Comeron et al. 2012). Understanding the genetic basis of this variability is essential for unraveling how and why recombination landscapes evolve (Ritz et al. 2017; Stapley et al. 2017; Johnston 2024) and the consequential impact on genome evolution.

Transposable elements (TEs), selfish genetic elements found in nearly all eukaryotic genomes, have emerged as potential drivers for the evolution of recombination landscapes. The ability of TEs to self-replicate and move between genomic locations not only influences diverse genome functions (e.g., (Gagnier et al. 2019; Cao et al. 2020; Lee et al. 2020; Payer et al. 2021)), but also generates substantial variation in the abundance, composition, and location of TE insertions between individuals (Cridland et al. 2013; Sudmant et al. 2015; Quadrana et al. 2016) and species (Wells and Feschotte 2020). Across taxa, TE abundance is widely observed to negatively correlate with recombination rates within a genome (Kent et al. 2017). While this association has largely been interpreted as a result of varying recombination rates leading to different effectiveness of selection purging TEs (Langley et al. 1988; Dolgin and Charlesworth 2008), accumulating observations suggest that TEs can instead be the cause rather than a consequence of a low recombination rate (Kent et al. 2017; Choi and Lee 2020). The potential suppressive impact of TEs on recombination is hinted by the widely observed low recombination rates in TE-infested pericentromeric heterochromatin (Kong et al. 2002; Ellermeier et al. 2010; Underwood et al. 2018; Fernandes et al. 2024). Within chromosome arms, clusters of TEs or high TE density are also less likely to co-occur with COs (Dooner and He 2008; He and Dooner 2009; Darrier et al. 2017) or DSBs (Pan et al. 2011; Yamada et al. 2017; Choi et al. 2018). On the other hand, the movement of TEs requires the formation of DSBs (Gasior et al. 2006), which aligns with the colocalization of CO hotspots within several TE families (Myers et al. 2005; Altemose et al. 2017), but contradicts the unaffected meiotic CO rates even in systems with highly elevated TE activities (Hemmer et al. 2020).

Several mechanisms could mediate the potential suppressive effects of TEs on COs. Polymorphic TEs are essentially insertions or deletions of several kilobases. They may have similar suppressive effects on COs as do other DNA structural variants, such as inversions and indels (Morgan et al. 2017; Crown et al. 2018; Rowan et al. 2019). In addition, as a consequence of host-directed epigenetic silencing to counteract the selfish replication of TEs, TEs within chromosome arms are oftentimes enriched with repressive epigenetic marks, in particular DNA methylation and H3K9me2/3 histone modification (Slotkin and Martienssen 2007). These repressive marks enriched at TEs could “spread” *in cis*, making TEs appear as islands of heterochromatin within the euchromatic genome (reviewed in (Choi and Lee 2020)). The very same repressive marks enriched in pericentromeric heterochromatin have been demonstrated to suppress the formation of DSBs (Peng and Karpen 2009; Choi et al. 2018) and the resolution of DSBs into COs (Maloisel and Rossignol 1998; Miller et al. 2012; Fernandes et al. 2024). While the repressive marks associated with TEs in the euchromatic genome are less extensive than those in pericentromeric heterochromatin (several kilobases *versus* several megabases), they may influence recombination through a similar mechanism. Consistent with this prediction, TEs that initially suppress DSB formation lose this inhibitory effect in mouse mutants lacking repressive epigenetic marks at TEs (Zamudio et al. 2015). Also, TE-containing motifs of CO hotspots in humans became CO coldspots upon acquiring epigenetic silencing marks (Altemose et al. 2017).

While previous studies suggest that TEs may suppress recombination, these effects were primarily inferred through comparisons of different genomic regions or by examining mutant backgrounds. Such approaches make it challenging to exclude potential confounding influences from local sequence context and to accurately assess the impact of TEs in natural populations. Moreover, many TEs identified as recombination modifiers are fragmented and likely inactive (Underwood and Choi 2019), and thus unlikely to contribute to the variation and evolution of recombination landscapes. In this study, we addressed these gaps by directly testing whether and how active, polymorphic TEs influence CO occurrence and drive varying recombination landscapes between individuals of the same species. We developed a novel approach using long-read PacBio sequencing on pools of recombinant offspring to efficiently, accurately, and cost-effectively identify COs in wild-type *Drosophila melanogaster* strains. With a few DNA extractions and PacBio SMRT cell sequencing, we identified nearly 3,000 COs and studied their distributions between *homologous* sequences with and without TE insertions in two *D. melanogaster* strains with distinct TE insertion profiles. In addition, we employed two other approaches—examining the distributions of COs in panels of recombinant inbred lines (RILs; (King et al. 2012b)) in relation to TE polymorphism and applying marker-assisted estimations of CO frequencies (Wei et al. 2020b) to isogenic strains differing in transgenically inserted TEs. These three orthogonal approaches collectively support the suppressive impact of TEs on CO occurrence, thereby contributing to the varying recombination landscapes observed between wild-type individuals of the same species.

## Results

### Proposed long-read pool-sequencing approach accurately identifies crossover

A typical approach to identify recombination events starts with crossing two inbred strains, followed by backcrossing the F1 to one of the parents and profiling genetic markers in F2 individuals (**Figure 1A**). The haplotype information of F2 offspring is critical for inferring CO events. Yet, the prevailing approach to acquiring that information is labor intensive and often cost-prohibitive, requiring separate DNA extraction and genotyping for every F2 individual. To circumvent this limitation, we proposed to sequence F2 offspring in pools using PacBio long-read sequencing, which enables efficient assaying of large numbers of individuals while preserving the local haplotype information (**Figure 1A**).

**Figure 1.**
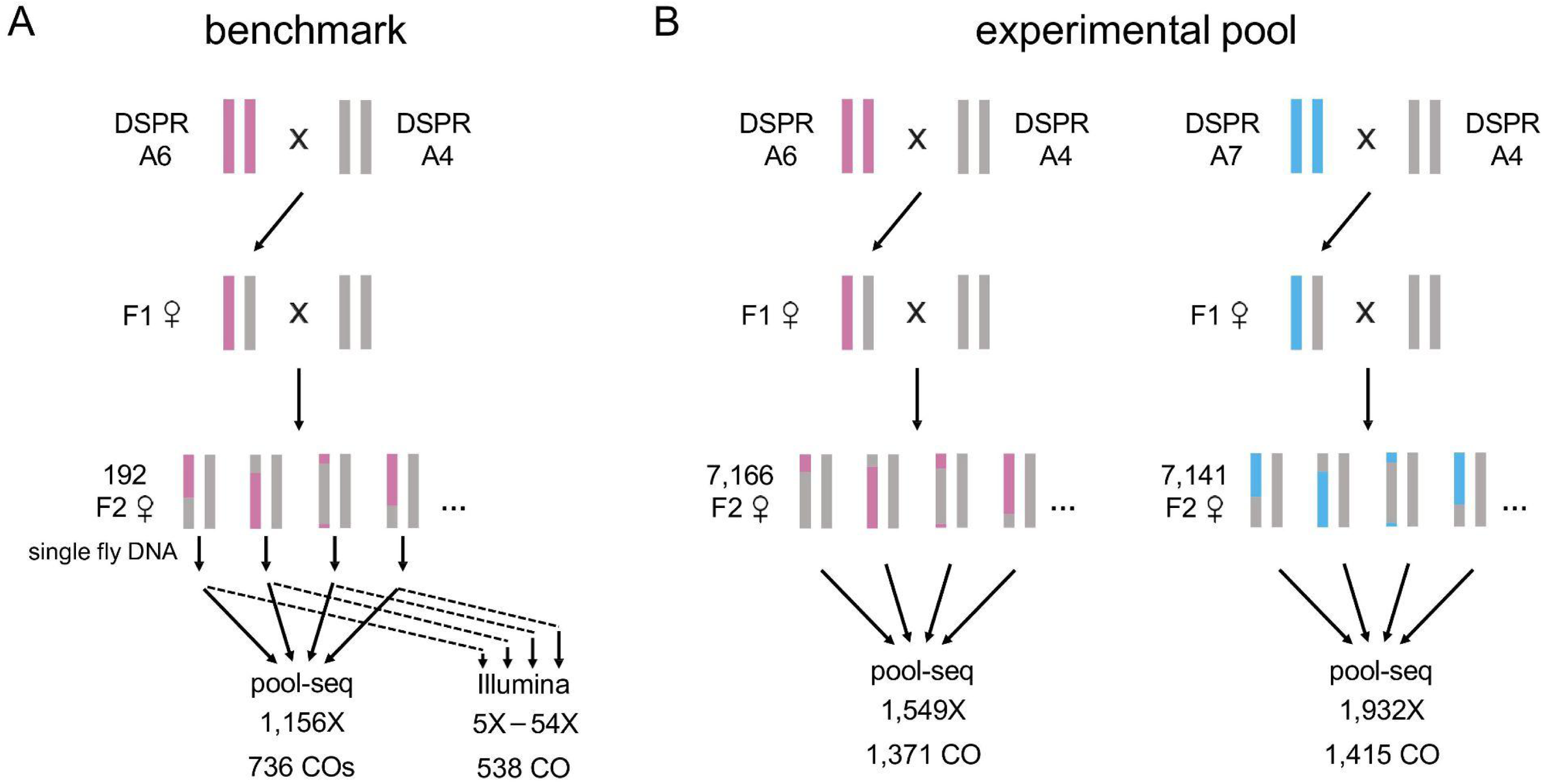
Experimental design of the benchmark data and experimental pools. (A) For the benchmark data, DNA was extracted individually from 192 female F2 backcrossed offspring of A6 x A4 crosses. Half of the DNA from each individual was sequenced by Illumina short-read sequencing, while the remaining DNA from all individuals was pooled in equal molar ratios and sequenced with PacBio CLR sequencing. (B) For the experimental pools, we set up two backcross experiments (A6 x A4 and A7 x A4). For each cross, over 7,000 female F2 backcrossed offspring were collected, pooled, DNA extracted en masse, and sequenced with PacBio CLR.

To investigate the feasibility of our approach, we generated a “benchmark” data set (**Figure 1B**). We performed crosses between two Drosophila Synthetic Population Resource (DSPR) founder strains (strains A4 and A6), which are highly inbred (King et al. 2012) and have been fully sequenced with PacBio long-read sequencing (Chakraborty et al. 2019). The short distance between SNPs (an average of 204bp) allows us to narrow down crossover events to within 1 kb (**Figure S1)**. We sequenced the DNA of 192 F2 female individuals both individually using Illumina short-read sequencing (short-read approach; with an average 11.5X depth of each individual) and as a pool using PacBio Continuous Long Read (CLR) sequencing (pool-seq approach; with a read depth of 1,156X; see Materials and Methods). COs were inferred by the transitions of SNP alleles from one parent to the other in the assembly (short-read approach) and in a read (pool-seq approach; see Materials and Methods).

We identified 538 CO events among 191 F2 individuals using short-read-based genome assemblies (one individual failed Illumina sequencing despite multiple attempts). From the same pool of 192 F2 individuals, our PacBio pool-seq approach detected 736 *reads* containing CO events. Of these, 688 (93.5%) overlapped with COs identified using the short-read method, indicating a low false-positive rate of 6.5% *per CO read* for the pool-seq approach. The sequencing depth per haplotype (3.01X) in the pool-seq method allowed for potential multiple-read capture of a single CO event. After merging COs within 100 bp, we identified 363 potentially unique CO events, further confirming a low false-positive rate of 8.3% *per CO event*.

Conversely, 203 of the 538 COs identified by the short-read approach were also detected using the pool-seq method. The COs not detected by our pool-seq approach (false negatives) were randomly distributed across the genome (**Figure S2**), indicating that such an approach shows no bias for or against particular genomic regions in CO detection. The relatively low recall rate of the pool-seq approach (37.7%) is likely due to our stringent CO calling criteria, which include removing reads with potential gene conversion events and requiring identified COs to be at least 2 kb away from alignment boundaries (see Materials and Methods). Relaxing these criteria significantly increases not only the recall rate, but also the false positive rate (**Table S1**). Therefore, for subsequent analyses, we maintained stringent criteria for calling COs in F2 pools to ensure high accuracy (see “Best practice to maximize the efficiency of CO detection with pool-seq approach” section for a discussion of the recall rate).

### Proposed long-read pool-seq method efficiently identifies large numbers of crossover

To test the predicted suppressive effects of TEs on recombination, we aimed to compare the distribution of COs between strains with different TE insertion profiles. We used three DSPR founder strains that have short average distances between SNPs (204 bp between A4 v.s. A6 and 217 bp between A4 v.s. A7) and conducted two backcross experiments (**Figure 1B**). For each cross, we pooled more than 7,000 F2 females and sequenced them in pool using PacBio CLR, with a final read depth of over 1,500x after filtering reads based on mapping quality and read length (**Figure 1B**, see Materials and Methods). In total, we identified 1,371 (A6 cross) and 1,415 (A7 cross) CO events, representing around 1,114 meiotic events based on the average number of 2.5 CO events per female genome per meiosis in *Drosophila* (Hughes et al. 2018). With the assumptions of equal sequencing coverage across the genome and the fact that half of the F2 DNA is from the nonrecombining parental chromosomes (**Figure 1B**), the expected number of detected COs should be 2.5 times half of the sequencing depth if our method is perfectly sensitive. Such a back-of-envelope calculation indicates an average of 65% recall rate for our proposed method (71% for A6 cross and 59% for A7 cross), which is much higher than that observed in the benchmark data (37.7%, see “Best practice to maximize the efficiency of CO detection with pool-seq approach” section). We compared the distribution of identified COs to the existing CO map generated by using Illumina short-read sequencing to genotype F2 individuals separately (Comeron et al. 2012) and found strong correlations between the two (*Spearman rank rho:* 53.7% (A6 cross), 50.6% (A7 cross), and 55.3% (averaged between the two crosses), *p* < 10^-16^, **Figure 2A** and **Figure S3**), which is in similar range to the reported correlations between short-read-based CO maps for any two *Drosophila* strains (∼45%, (Comeron et al. 2012)). This observation suggests that our proposed approach effectively and efficiently recapitulates the genome-wide variation in COs.

**Figure 2.**
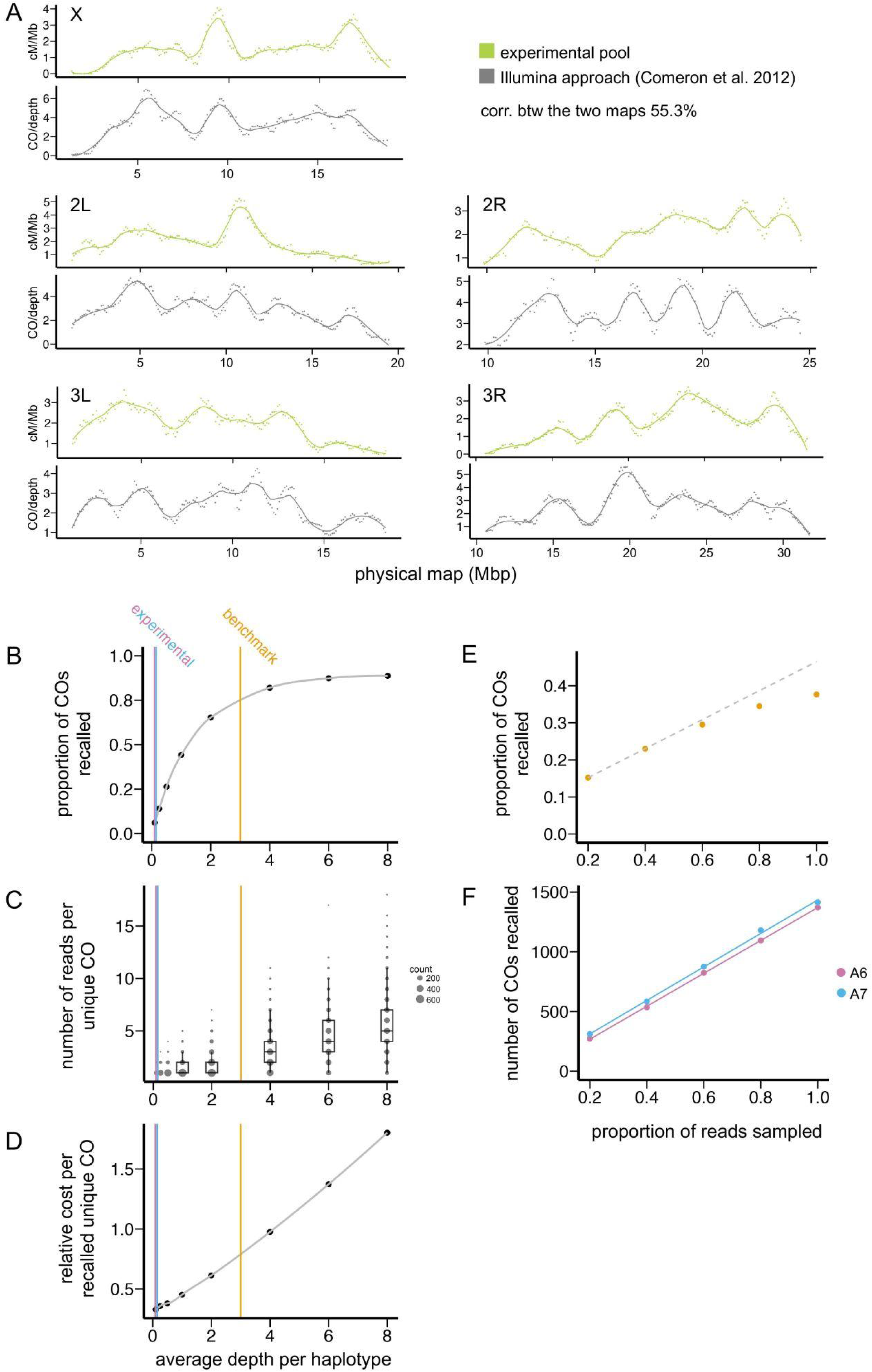
CO maps inferred by the proposed pool-seq approach and the impact of sequencing depths per haplotype on the sensitivity and efficiency of the approach. (A) The read-depth normalized CO numbers identified in our experimental pools (in every 100 windows and combining the two strains) for each chromosome is shown on the top, and a previously generated recombination map based on a short-read Illumina sequencing approach (replotting from **(Comeron et al. 2012)**) is showing on the bottom. The distribution of COs was smoothed by a sliding window of 1 Mb with a step size of 100 kb and LOESS with a 20% span. See **Figure S3** for the distribution of CO numbers of individual strains. (B-D) The recall rate of COs for different depth-per-haplotype ratios using pool-seq simulations. As the sequencing depth per haplotype increases, the proportion of COs recalled increases (B), as well as the average number of reads recovering the same CO event (C). When more than one read is recovering the same CO event, a CO is called repeatedly, leading to high sequencing cost per uniquely called CO (D). Here, the sequencing cost is estimated as the read depth divided by the number of CO recalled in a simulation pool. The observed depth-per-haplotype ratios for the benchmark data (orange) and experimental pools (blue and pink) are denoted as vertical lines. (E) For the benchmark data, the CO recall rate of the pool-seq approach was estimated by calculating the proportion of COs identified by the short-read method that were also detected by the pool-seq approach at a given sequencing coverage. The CO recall rate plateaus as the read depth increases, which is likely due to the increased sequencing depth per haplotype (Figure 2C). (F) For the experimental pool, the number of COs recalled under different sequencing depths was inferred by downsampling. The nearly linear relationship suggests the efficient utilization of the sequencing efforts to detect COs in the experimental pool.

### Potential biases of the proposed long-read pool-seq approach in detecting crossover

To ensure that any observed association between CO occurrence and TE presence is not due to detection biases, we conducted in silico simulations to investigate how potential confounding factors may influence CO detection. This section focuses on these simulations, while subsequent sections discuss analyses of this issue using experimental data. We first examined how the relatively high error rate of PacBio CLR reads might influence CO detection by simulating reads both without (to assess false positives) and with (to assess recall rate) CO events (see Materials and Methods). Our analysis found no false CO detections when reads were simulated from non-recombinant parental genomes, indicating high precision in our approach despite high sequencing error rates. For simulations with CO events, our approach achieved an 81.4% recall rate for reads that are properly mapped, overlap with CO events, and pass our stringent filtering criteria (recall rate *per read*). Relaxation of these filtering criteria further increased the recall rate (**Table S2**). The inability to achieve a 100% recall rate is likely partially due to some false negative events occurring in genomic regions with few or no SNPs between the two parents (**Figure S4**). However, this issue should not affect the analysis of our benchmark data, as COs in such low-diversity regions would not have been detected by the short-read approach either.

Next, we assessed whether the presence of TEs could influence the detection of COs. Surprisingly, simulated reads that span TEs identify COs at a significantly higher rate than reads aligned to other genomic regions without TEs (70% *v.s.* 57% detection rate, *Fisher’s Exact Test, p* < 10^-16^). This bias should make any observed negative association between the occurrence of CO and the presence of TEs conservative. Interestingly, we detect a similar effect for reads spanning other non-TE structural variants (71% (with SV) *v.s.* 58% (without SV), *Fisher’s Exact Test, p* < 10^-16^). To investigate the impacts of sequencing depth, read length, and SNP density on detecting CO, we correlated these factors with CO numbers detected with simulated PacBio CLR data in windows that are *far from* TEs (see Materials and Methods). We found the number of detected CO positively correlated with sequencing depth (*Spearman rank rho* = 0.03, *p* = 0.04) and SNP density across windows (*Spearman rank rho* = 0.06, *p* =10^-5^). On the other hand, there is no such association between read length and the number of CO detected (*Spearman rank correlation test, p* = 0.49). These observations suggest a need to consider sequencing depth and SNP density when investigating the associations between the distributions of COs and TEs, which we implemented below.

### Best practice to maximize the efficiency of CO detection with pool-seq approach

The significant difference in recall rates between our benchmark data (37.7%) and experimental pool (65%) raises important questions about the factors causing these disparities and cost-effective approaches for implementing our proposed pool-seq method. Our simulation results indicate that both sequencing errors and stringent CO-calling criteria lead to failures in detecting COs (see above, **Table S2**). Additionally, the unequal contribution of DNA from F2 individuals to the final sequencing pool could further lower the recall rate. This is evidenced by a larger number of COs being detected by more than the expected number of reads in the benchmark pool (**Figure S5**). In contrast, few COs were detected by multiple reads in the experimental pool, suggesting minimal impact from unequal pooling in this case. Unlike the benchmark data, where extracted DNA was pooled post hoc and may be sensitive to measurement errors, the DNA of the experimental pool was extracted en masse from a large number of F2 individuals, which is more likely to achieve random sampling across individuals and reduce bias in representation.

Another major difference between the benchmark data and the experimental pool is the ratio of sequencing depth per recombinant F2 haplotype (referred to as depth per haplotype hereafter; 3X for the benchmark data *versus* 0.12X for the experimental pool). We used additional simulations to explore how this depth-per-haplotype ratio may influence recall rates (see Materials and Methods). As the depth-per-haplotype ratio increases, the recall rate steadily improves, plateauing at approximately 85% (**Figure 2B**). The recall rate never reaches 100%, likely due to the influence of sequencing errors and SNP availability. Interestingly, the number of simulated reads recovering *the same* CO event also increases (**Figure 2C**). Such an observation suggests that many of the sequencing reads could be “wasted” in identifying new COs when the depth-per-haplotype ratio is large (**Figure 2D**). Consistently, through subsampling reads, we found a nearly perfect linear relationship between sequencing depth and the number of CO identified in the experimental pool (**Figure 2E**), but a plateau curve for the benchmark data (**Figure 2F**), illustrating how the sequencing depth per haplotype influences the cost effectiveness of recovering COs.

Given the random distribution of false negative events across the genome (**Figure S2**), we reasoned that, when implementing our pool-seq approach, the primary goal should be to maximize the number of COs detected *per unit of sequencing depth*, rather than attempting to recover all COs in the sequenced pool. When the depth-per-haplotype ratio in the sequencing pool is low, the probability of COs being captured by multiple reads minimizes (**Figure 2C**), which avoids potential confoundment in the downstream analysis (e.g., distinguish between independent nearby CO *versus* repeated CO capturing) and the waste of sequencing costs (**Figure 2D**). Accordingly, we recommend the following best practice: extract DNA en masse from a large number of individuals and incrementally increase sequencing depth while maintaining a low depth-per-haplotype ratio until the desired number of COs are recovered. This strategy balances between sequencing costs and CO detection efficiency.

### CO occurrence is suppressed around euchromatic TEs when compared to other euchromatic regions

We first investigated the predicted negative impact of TEs on CO occurrence by comparing *euchromatic* windows with and without TEs within the individual strain. We estimated the number of COs in 5 kb windows upstream and downstream to TEs (TE flanking windows) and in 10 kb windows that are at least 30 kb away from TEs (control windows, **Figure 3A**). TE flanking windows tend to have lower sequencing depth and fewer SNPs than the control windows (*Mann-Whitney U test, p <* 0.05 for all comparisons; **Figure S6**). These differences could potentially confound our analysis because, similar to observations with simulated data (see above), the number of identified COs across non-TE windows are positively correlated with sequencing depth (*Spearman rank rho* = 0.018, *p* = 0.12 (A6) and *rho* = 0.055, *p* < 10^-6^ (A7)) and SNP numbers (*Spearman rank rho* = 0.095, *p* = 6.0×10^-10^ (A6) and *rho* = 0.10, *p =* 6.7×10^-12^). Accordingly, we included sequencing depth and the number of SNPs as co-variables when assaying the impacts of TEs on the distributions of COs. On the other hand, read length would not have confounded our analysis because we found no associations between that and the number of COs detected (*Spearman rank correlation tests, p* = 0.15 (A6) and 0.14 (A7)).

**Figure 3.**
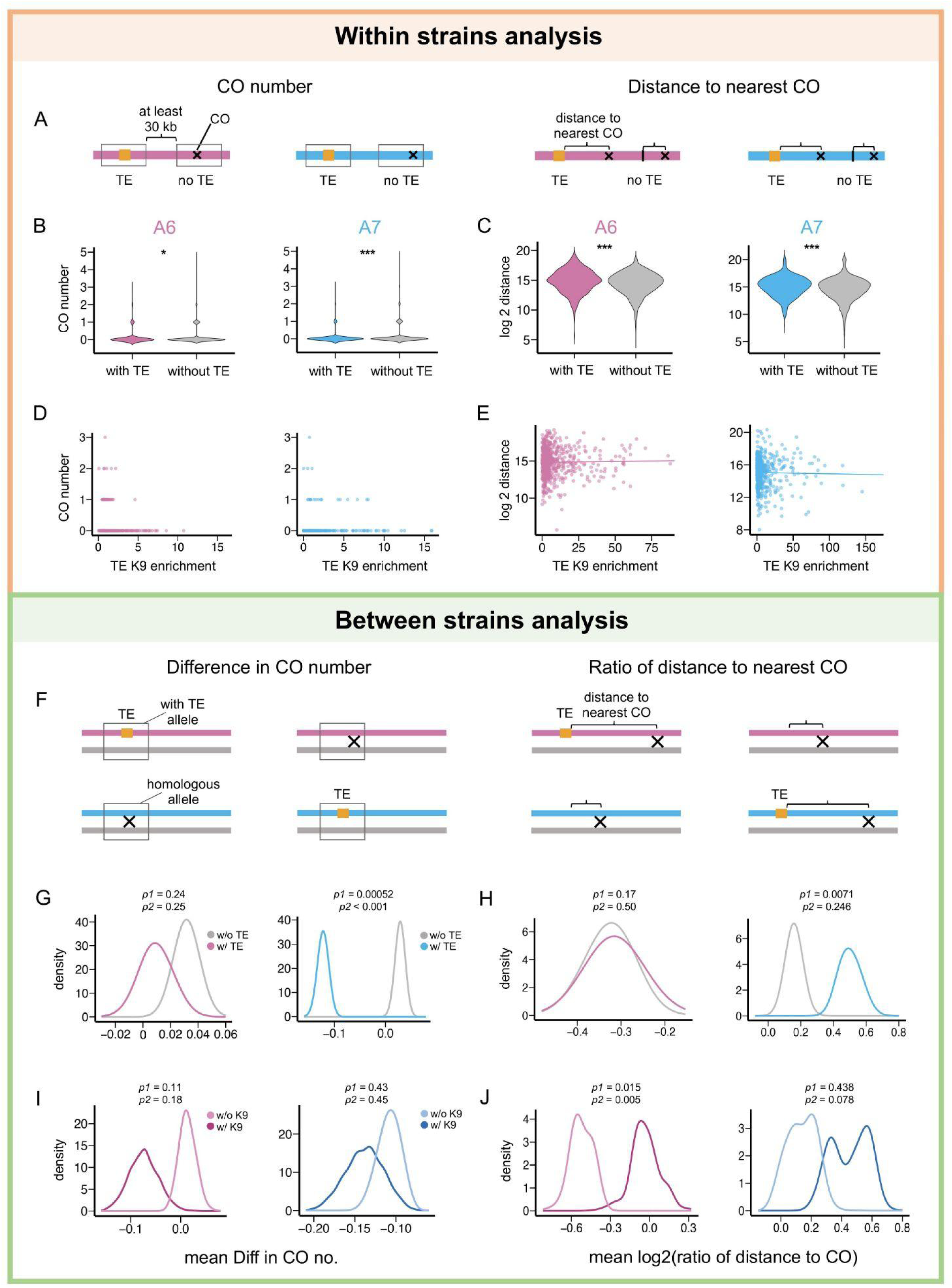
Euchromatic TEs suppress CO occurrence. (A-E) Within-strain analysis to investigate the impact of TEs on the occurrence of COs. (A) Cartoon illustrating measures for the occurrence of COs: CO numbers of 5 kb windows flanking focal sites and the physical distance from the focal site to the nearest CO. (B) Within-strain comparisons of CO numbers for TE windows (451 and 340 windows for A6 and A7, respectively) and control windows without TEs (4,185 and 4,720 windows for A6 and A7 respectively). (C) Within-strain comparisons of the distance to the nearest COs TEs (624 and 505 TEs for A6 and A7, respectively) and controls (4,256 and 4,924 control sites for A6 and A7, respectively). (D and E) Associations between TE-induced H3K9me3 enrichment and CO numbers (D) and the distance to nearest CO (E). (F-J) Between strain analysis to test the impact of TEs on the occurrence of COs. (F) Cartoon illustrating measures for the impacts of TEs on CO occurrence: difference in CO numbers in the TE flanking windows (left) and log2 ratio of the distance to nearest CO (right) between homologous alleles with and without TEs. (G) Shown distributions compare the mean differences in CO numbers for 1,000 downsampled sets of TE pairs that have equal sequencing coverage between homologous alleles with and without TEs (345 and 232 TEs for A6 and A7, respectively) and the same number of downsampled control window pairs as TEs (bootstrapped from 1,434 and 1,507 control windows in A6 and A7, respectively). Because most windows overlap with no COs, the distribution of the median difference in CO number is less visually informative than the mean presented here. (H) Shown distributions compare the mean differences in distance to the nearest COs for 1,000 downsampled sets of TE pairs (489 and 360 TEs in A6 and A7, respectively) and the same number of control windows as TEs (bootstrapped from 1,831 and 2,041 control windows in A6 and A7, respectively). (I) Comparisons for the difference in CO numbers between TE pairs with (70 and 61 TEs in A6 and A7, respectively) and without H3K9me3 enrichment (135 and 90 TEs in A6 and A7, respectively). (J) Comparisons for the ratios of the distance to nearest CO between TE pairs with high and low H3K9me3 mass (123 and 93 TEs in either category for A6 and A7, respectively). A significant level was obtained via both the *Mann-Whitney U* test (*p*1) and the bootstrapping test (*p*2) for between-strain analysis (see main text). The numbers of TEs in (B, G) differ from those of (C, H) because, for CO number, we required at least five SNPs within both 5 kb flanking windows (B, G), while for analysis using distance to nearest CO (C, H), this was not criterion was not required. The smaller number of TEs studied in (I and J) analysis when compared to (G and H) is due to the stringent filtering of TE pairs whose without TE alleles also show some enrichment of H3K9me3. w/ TE: TE windows/sites; w/o TE: control windows/sites without TEs; w/ K9: with H3K9me3 enrichment; w/o K9: without H3K9me3 enrichment. For (B-E) *Mann-Whitney U* test, \**p* < 0.05; \*\*\**p* < 0.001.

For both strains, we found that TE flanking windows have significantly lower numbers of CO than control windows (*Mann-Whitney U test, p* = 0.017 (A6) and 1.5×10^-4^ (A7); **Figure 3B**), a result that is robust when controlling for the confounding effects of sequencing depth and number of SNPs using generalized linear models (regression coefficients of TE effect: −0.20, *p* = 0.24 (A6) and −0.71, *p =* 0.00083 (A7); see Materials and Methods). In addition, we compared the physical distance from the focal site to the nearest CO events, which should be longer when CO occurrence is suppressed around the focal site. Consistent with analysis comparing CO numbers, the distance from TE insertion site to the nearest CO is significantly longer than that of the control site in both strains (*Mann-Whitney U test, p* = 8.6×10^-7^ (A6) and *p* = 6.9×10^-7^ (A7); Regression coefficient for TE effect: 1.52 (A6) and 1.29 (A7), *p* < 10^-16^ for both; **Figure 3C**). These observations support the prediction that TEs suppress the occurrence of COs.

We next investigated what biological attributes make a TE exert stronger suppressive effects on COs. While we did not find differences in CO numbers around different classes of TEs (RNA *versus* DNA, and LTR *versus* non-LTR *versus* TIR, regression coefficient *p* > 0.05 for all pairwise comparisons; **Table S3**), RNA-based TEs, when analyzed jointly or separately, lead to significantly longer distances to the nearest CO than DNA-based TEs in one strain (A6; Regression coefficient: 0.506 (RNA *versus* DNA), 0.508 (LTR *versus* DNA) and 0.505 (non-LTR *versus* DNA), *p* < 0.01; **Table S3**). We then tested whether the enrichment of H3K9me3 may act as a driver for the suppressive effects of TEs on COs. Two measures for the enrichment of H3K9me3 were used: the average H3K9me3 enrichment level in TE flanking windows (adjacent H3K9me3 enrichment) and accumulated H3K9me3 across the entire extent of H3K9me3 spread from TEs (H3K9me3 mass; **Figure S7**). We used the former measure for analysis using CO numbers and the latter for analyzing distance to the nearest CO, because the extent of the suppressive effect of H3K9me3 is unknown. We found significant negative associations between adjacent H3K9me3 enrichment and the number of COs identified in one strain (A6: *Spearman rank rho* = −0.097, *p =* 0.028; regression coefficient of H3K9me3 enrichment −0.42, *p* = 0.047; **Figure 3D**), while associations using H3K9me3 mass were not significant (**Figure 3E; Table S3**). Similar to previous studies, we also found differences in the level of H3K9me3 enrichment between TE classes (Lee and Karpen 2017; Huang et al. 2022) (**Table S4**). Alongside our finding that RNA-based TEs exert stronger suppressive effects on CO occurrence, we sought to differentiate whether the observed negative association between H3K9me3 enrichment and CO occurrence is due to TE class or the enrichment of H3K9me3 itself. We repeated the above analysis *within TE class,* and still found significantly negative correlations between the enrichment of H3K9me3 and CO number for RNA-based TEs (A6: *Spearman rank rho* = −0.18, *p* = 0.0014 (RNA-based TE), *rho* = −0.18, *p =* 0.015 (LTR), *rho =* −0.18, *p =* 0.057 (non-LTR); regression coefficient of H3K9me3 enrichment −0.59, *p* = 0.057 (RNA-based TE), −0.46, *p* = 0.12 (LTR), and −1.90, *p* = 0.053 (non-LTR); all other comparisons, including analysis using distance to CO, have *p* > 0.05, see **Table S3** for details), confirming the impact of TE-induced enrichment of repressive marks on CO occurrence. Interestingly, TEs *without* H3K9me3 also have fewer CO numbers (regression coefficient of TE presence −0.086, *p* = 0.68 (A6) and −0.684, *p =* 0.031 (A7)) and show longer distance to the nearest CO (regression coefficient of TE presence 1.38 (A6) and 1.44 (A7), *p* < 10^-16^ for both) than the control windows. These TEs may impact CO occurrence similar to other SVs, an effect expected to be stronger for longer TEs. We tested this possibility by focusing on TEs *without* the enrichment of H3K9me3, but found no significant associations between TE length and the number of COs detected nor distance to nearest COs (Spearman rank correlation analysis and regression analysis, *p* > 0.05 for all comparisons, **Table S3**). In summary, our observations suggest that TEs, especially RNA-based TEs and TEs with high local enrichment of repressive marks, suppress the occurrence of COs in their vicinity.

### CO occurrence is suppressed in the presence of euchromatic TEs when compared to homologous sequences without TEs

While our above analysis found a reduced occurrence of COs around TEs when compared to TE-free genomic regions within a strain, similar patterns could also be driven by other local features influencing CO occurrence, as well as TE insertion preference or differential selection against TEs (reviewed in (Kent et al. 2017)). To investigate the suppressive impacts of TEs on CO while accounting for these potential confounding effects, we estimated the difference in two indexes, the number of COs within windows and the distance to the nearest CO, between *homologous alleles* with and without TEs (TE set, **Figure 3F**). These estimates were compared to those of control sites at least 30 kb away from TEs (control set) to account for the difference in recall rate and, potentially, genetic map length between the two strains (see Materials and Methods). Because of the pronounced impacts of sequencing depth on the detection of COs (see above), we inferred indexes of CO distribution after downsampling sequencing reads to ensure equal sequencing coverage between homologous alleles. In addition, our analysis controlled for SNP number (see Materials and Methods) because the ratio of the number of SNPs between homologous alleles is marginally insignificantly different between TE sites and control sites (*Mann-Whitney U test, p* = 0.06 (A6) and 0.08 (A7)), Significance levels of differences in CO distribution between homologous alleles with and without TEs were determined by two approaches: (1) the *median Mann-Whiteney U test p-values* for the comparisons between the TE set and control set across 1,000 downsampled data and (2) comparing the observed difference in CO indexes of the TE set to the null distribution generated by bootstrapping control sets (see Materials and Methods).

We first tested whether the presence of TEs negatively impacts CO occurrence, which should be reflected as smaller values for the difference in CO number but larger values for the ratios of distance to the nearest CO for the TE set than those of the control set (**Figure 3F**). Consistently, compared to the control set, the TE set has a significantly lower difference in CO number (*Mann-Whitney U test, median p:* 0.24 (A6) and 0.00052 (A7); *bootstrapping p:* 0.25 (A6) and < 0.001 (A7), **Figure 3G and Figure S8**) and larger ratios of distance to CO (*Mann-Whitney U test, median p:* 0.17 (A6) and 0.0071 (A7); *bootstrapping p:* 0.501 (A6) and 0.246 (A7), **Figure 3H and Figure S8**). We found limited differences in the suppressive impact of TEs of different classes and types, with non-LTR showing significantly larger ratios in the distance to COs than TIRs in one of the two strains (*Mann-Whitney U test, median p* = 0.037, all other comparisons *p* > 0.05, see **Table S5**).

To investigate whether the enrichment of H3K9me3 around TEs contributes to varying suppressive effects of TEs on COs, we compared the differences in CO numbers between TEs with and without H3K9me3 enrichment in their 5kb flanking window (adjacent enrichment level > 1). Notably, non-TE induced enrichment of repressive marks does occur in the euchromatic genomic region (e.g., (Riddle et al. 2011)) and, to mitigate the consequential confounding effect, we excluded windows where alleles without TEs similarly show enrichment of H3K9me3. TEs with H3K9me3 enrichment tend to have lower differences in CO number than TEs without, although the difference is not statistically significant (*Mann-Whitney U test, median p:* 0.11 (A6) and 0.43 (A7); *bootstrapping p:* 0.18 (A6) and 0.45 (A7), **Figure 3I and Figure S9**).

Alternatively, we correlated the level of H3K9me3 enrichment to the difference in CO number and found a significant negative association for one of the two strains (*median Spearman correlation coefficient*: −0.12 (A6, median *p* = 0.043) and −0.03 (A7, median *p* = 0.36), **Figure S10**). Removing four *Roo* elements near COs, despite being highly enriched with H3K9me3, in the insignificant strain leads to a stronger correlation (*median Spearman correlation coefficient*: −0.10, median *p* = 0.11 in A7). We also performed a similar analysis using ratios of the distance to the nearest CO and used the total H3K9me3 mass to categorize TEs (high and low H3K9me3 enrichment), excluding those with enrichment for H3K9me3 in the TE-free homologous alleles (see Materials and Methods). TEs with high H3K9me3 mass show significantly larger ratios of the distance to the nearest CO, especially in one of the two strains (*Mann-Whitney U test, median p:* 0.015 (A6) and 0.438 (A7); *bootstrapping p:* 0.005 (A6) and 0.078 (A7), **Figure 3J and Figure S9**). In the same strain, we also found a significant positive correlation between H3K9me3 mass and the ratios of the distance to the nearest CO (*median Spearman correlation coefficient*: 0.11, *p* = 0.041 (A6); **Figure S10**). Importantly, significant negative associations between TE-induced enrichment of H3K9me3 and difference in CO numbers remain when restricting the analysis to RNA-based TEs (*median Spearman correlation coefficient*: −0.24, *p* = 0.003 (RNA-based TEs), −0.26, *p =* 0.005 (LTR)), −0.24, *p* = 0.1 (non-LTR); see **Table S5** for all other comparisons), suggesting such an association is not due to difference in H3K9me3 enrichment between TE classes (**Table S4**). Finally, similar to *within-strain* analysis, we found that even TEs *without* H3K9me3 show suppressive effects on COs in one of the strains (CO number: *Mann-Whitney U test median p =* 0.030 and bootstrapping *p =* 0.016 (A7); **Table S5**), but found no correlations between TE length and difference in CO distribution for TE insertions where alleles with and without TEs are both depleted of H3K9me3 enrichment (*p* > 0.05 for all comparisons, **Figure S11, Table S5**). Overall, our between-strain analyses that compare the distribution of COs between homologous alleles with and without TEs corroborate our findings observed within strains: TEs, especially those that lead to local enrichment of repressive marks, suppress the occurrence of COs in the euchromatic genome.

### Suppressive effect of TEs on recombination is supported by orthogonal approaches

To further corroborate our observed suppressive effects of TEs on COs in the euchromatic genome, we used two orthogonal approaches to independently test this finding. In the first approach, we investigated the distributions of CO breakpoints in the RIL panel of DSPRs **(King et al. 2012b)** with respect to the presence/absence of TE insertions. We inferred the CO breakpoints in the subset of Illumina-sequenced RILs, estimated the distance between an euchromatic TE insertion site to the nearest recombination breakpoint, and compared such distances between homologous alleles with and without TEs (see Materials and Methods). Consistently, alleles with TEs have a significantly longer physical distance to the nearest CO breakpoints than homologous alleles without TEs (**Figure 4A and 4B**; *paired Mann-Whitney U test, p* < 10^-16^), with the presence of TEs adding a median of 207,894bp to the nearest breakpoints. Analysis focusing on insertion sites where the median distance to breakpoint is within 100kb reaches the same conclusion (presence of TEs add 230,454bp, *paired Mann-Whitney U test, p* < 10^-16^). Among all TE insertion sites studied, 66.82% of them have longer physical distances to the nearest CO breakpoint when the TE is present than when the TE is absent, a proportion that is significantly larger than randomly sampled genomic locations with matching chromosomal distributions and numbers (**Figure 4C**, permutation test *p* < 0.001). However, we did not find differences in the suppressive effects of TEs of different classes (*Mann-Whitney U test, p >* 0.05 for all comparisons).

**Figure 4.**
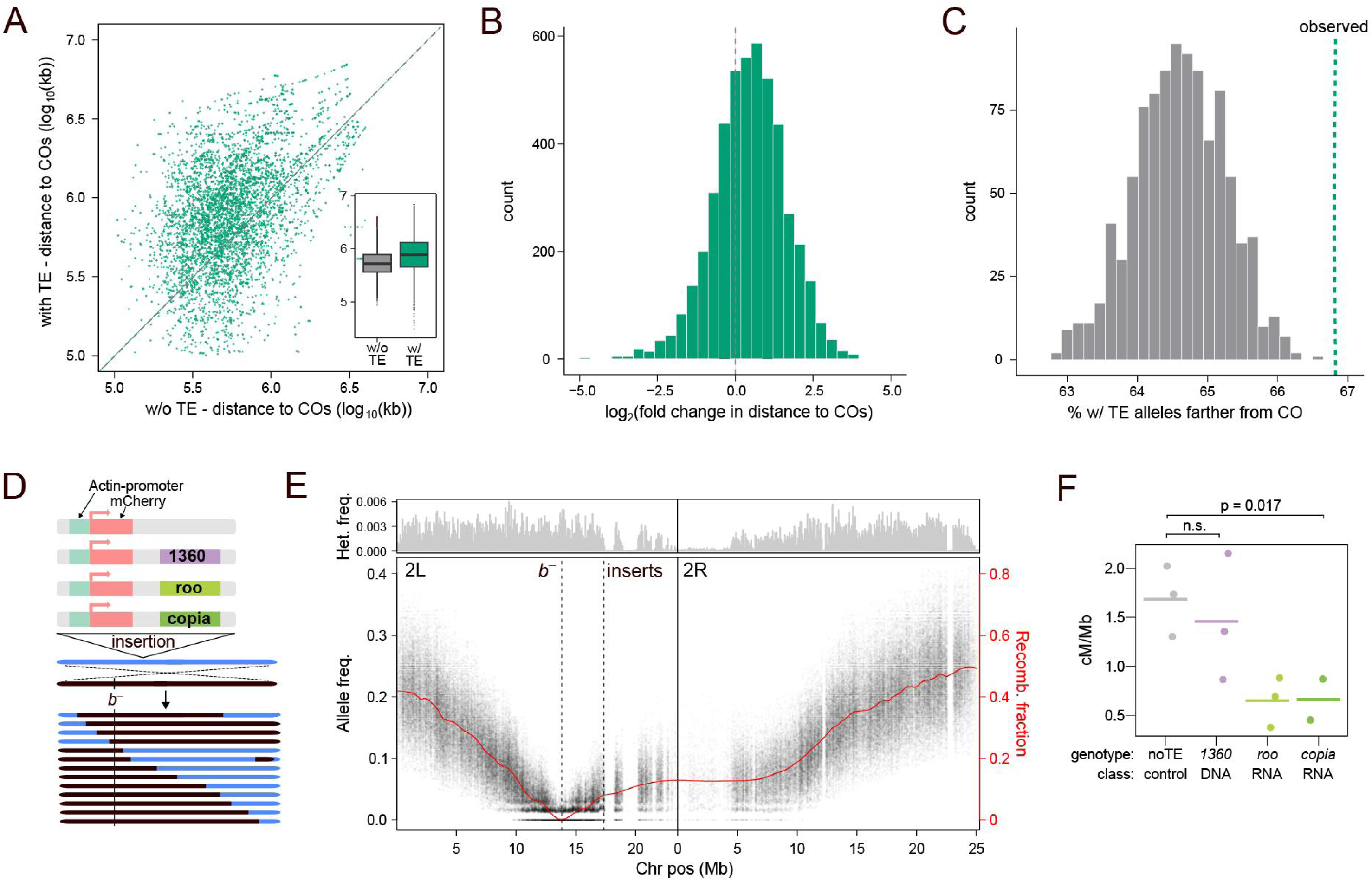
Orthogonal approaches support the conclusion that TEs suppress CO occurrence. (A-C) Comparisons of the distance to nearest CO between homologous alleles with and without TEs using RILs of DSPR. (A) log 10 distance to the nearest CO breakpoints from TE insertion sites between alleles without (w/o, x-axis) and with (y-axis) with TEs, which is biased towards larger distance for with TE alleles. Insert: boxplot showcasing the differences in distance to the nearest breakpoint between homologous alleles with and without TEs. (B) The distribution of fold change in the distance to the nearest CO breakpoints from TE insertion sites between homologous alleles with and without TEs. (C) Comparisons of the observed proportion of TE insertions with (w/) TE alleles having a farther distance to the nearest CO breakpoints than the homologous alleles (dashed green line) to the distribution of that of randomly sampled genome locations with matching properties (gray). (D) Cartoon representation of the transgenic strains used and how genetic distance is captured in allele frequency of marker selected pool. (E) After sequencing, allele frequency is estimated at every diagnostic SNP site where the parental strains are fixed for different alleles. The density of such sites is displayed in the top panel (in 100kb windows). Decay of allele frequency around *black*, is then used to estimate the recombinant fraction, which approximates the genetic distance (red). The rate of change (slope) then approximates the recombination rate. (F) Estimated recombination rate using SNPs ±500kb of the selected locus *black* in crosses with different transgenically introduced TEs. Significance differences between different TE insertions were estimated using a linear model: CO rates ∼ class + genotype. Because the genotype effect was not significant (*p* = 0.98), only the class effect was analyzed using the ANOVA and TukeyHSD tests.

In the other approach to test the causal impacts of TEs on recombination, we compared the distribution of COs around alleles with and without transgenically introduced TEs in a controlled genetic background. Such an approach mitigates the potential difference in genetic background, which is hard to ascertain in the above genome-wide approaches. Specifically, we inserted mCherry containing transgenic constructs with different TE sequences into the same euchromatic location (see Materials and Methods). To assess local changes in recombination rate in the presence of TE sequences, we employed a recently developed technique that infers chromosome-wide recombination rates from bulk sequencing of marker-selected pools (Wei et al. 2020b). We chose to integrate two RNA-based TEs that showed the strongest *cis* spreading of repressive marks in our previous study (*roo* and *copia,* (Lee and Karpen 2017; Huang et al. 2022)), and a DNA-based TE identified to exert *cis* epigenetic effects (*1360,* (Sentmanat and Elgin 2012)). In a F2 recombinant backcross (**Figure 4D**), we selected F2 homozygotes with the visible *black* marker, which is ∼7 Mb from the transgenic insertion site. In the selected F2 pool, the allele frequency of the inserted chromosome (blue in **Figure 4D**) will increase as a function of increased genetic distance from *black* (**Figure 4E**), and we extrapolated the local recombination rate as the rate of change in genetic distance after accounting for paternal contribution and secondary fitness effects. Within the 500kb flanking windows of different transgenic constructs, there is significant variation in the recombination rate (ANOVA F = 7.2, *p* = 0.017), with RNA-based TEs being significantly lower than the control (TukeyHSD test, adjusted *p* = 0.02; **Figure 4F**). It is worth noting that the transgenic background coincidentally shares several large haplotype blocks just downstream of the transgenic insertion site, leading to a sharp decrease in informative SNP sites, and, therefore, suboptimal estimation of allele frequency. Nevertheless, both approaches provide orthogonal support for the suppressive effects of TEs on local recombination rate.

## Discussion

By developing a new long-read based method to identify COs among pools of recombinant individuals, we compared the distribution of ∼2,800 COs with respect to TE insertions. Our analysis revealed that TEs generally suppress CO occurrence, especially for RNA-based TEs and TEs with enrichment of repressive epigenetic marks. After normalizing the varying detection rates of COs between the two *D. melanogaster* strains studied, we found the presence of TEs is associated with a 28% reduction in CO occurrence, with TEs leading to H3K9me3 enrichment having an even larger effect (41% reduction). Importantly, our orthogonal approaches for comparing CO frequencies between homologous alleles with and without TEs reached the same conclusion, further corroborating the suppressive effects of TEs on recombination. Importantly, all the TEs included in our study are polymorphic and likely still active. This is in contrast to TEs suggested to be recombination hotspots in plants and mammals, where most of those TEs are no longer active and oftentimes share similar genetic features (e.g., DNA motifs) with other non-TE recombination hotspots (Myers et al. 2005; Altemose et al. 2017; Underwood and Choi 2019).

Our method of applying PacBio long-read sequencing on pooled recombinant F2 is similar in concept to previous work using linked reads to detect COs in gametic pools (Dréau et al. 2019; Sun et al. 2019) but with higher accuracy (false positive rate 6.5% *v.s.* 14.9%), easier analysis, and greater cost-effectiveness. Our algorithm identifies COs by detecting the transition of SNPs from one parent to the other within a single PacBio read, eliminating the need to reconstruct sequencing molecules and filter shared barcodes as required with linked-read sequencing (e.g., (Dréau et al. 2019; Xu et al. 2020)). For individual *Drosophila* strain, our approach can detect COs of over five hundred meiosis events with just one DNA extraction, one PacBio library preparation, and two PacBio SMRT cell sequencing. The entire process cost roughly $4,000 several years ago and is likely much cheaper now. In contrast, identifying a similar number of COs using the short-read sequencing approach would require over five hundred separate DNA extractions and library preparations, more than doubling the costs in reagents and sequencing alone while also demanding significantly more labor and time.

Importantly, our benchmark data and in sillico simulations not only allow us to estimate the accuracy of the approach, but also to identify potential sources of false negative events and inform practices to maximize the efficiency of our approach. Our analyses reveal that, while a high sequencing depth per F2 recombinant haplotype leads to improved CO recall, it also results in redundant capture of the same COs, an inefficient use of sequencing efforts (Figure 2B-D). In addition, such a high depth-to-haplotype ratio is susceptible to unequal sampling of F2 individuals. Considering the randomly distributed false negative events in the benchmark data (**Figure S2**) and the nearly linear relationship between sequencing depth and the number of unique COs detected in the experimental pools (Figure 2F), we recommend extracting DNA en mass from thousands of F2 recombinant individuals and sequencing the DNA in batches until the desired number of COs is detected. Crucially, a low sequencing depth-to-haplotype ratio should be maintained to help maximize the number of COs detected per unit of sequencing effort. It is worth noting that we were unable to infer gene conversion events accurately due to the relatively high error rate of PacBio CLR (∼10%, (Logsdon et al. 2020)), the distance between SNPs in the parental strains, and the typically short track length of gene conversion events (∼500bp, (Comeron et al. 2012; Miller et al. 2012)). Recent applications of the much more accurate, but shorter, PacBio HiFi sequencing demonstrate the possibility of identifying other recombination products from gametic pools (Porsborg et al. 2024; Schweiger et al. 2024). Further benchmarking, like those developed in this study, will help assess the specificity and sensitivity of using PacBio HiFi sequencing to detect various recombination products.

Our findings reveal that RNA-based TEs are more likely to suppress CO occurrence. This finding aligns with previously reported heterogeneity in the effects of different TE classes on recombination, particularly the more pronounced negative associations of RNA-based TEs with the formation of DSBs (Yamada et al. 2017; Choi et al. 2018) and occurrence of CO (Myers et al. 2005; Darrier et al. 2017). However, RNA-based TEs tend to result in stronger local enrichment of repressive marks in *Drosophila* (Huang et al. 2022). Their negative effects on COs might be mediated by the known suppressive impacts of these marks on the various steps of COs (Maloisel and Rossignol 1998; Peng and Karpen 2009; Choi et al. 2018), which is supported by our observed negative associations between the H3K9me3 enrichment around TEs and CO occurrence *within* TEs classes. Interestingly, TEs without the enrichment of H3K9me3 also significantly associate with reduced CO occurrence in one of the two strains, indicating that other mechanisms also mediate the suppressive effects. One possibility is that, similar to the reported effects other structural variants (Borts and Haber 1987; Morgan et al. 2017), sequence heterogeneity introduced by TEs interferes with homolog pairing, leading to the resolution of DSBs into non-CO products. Alternatively, because our approach can only detect COs in surviving adults, COs that reduce an individual’s survival are likely underrepresented in the F2 pools. This bias is particularly likely when non-homologous TE sequences serve as templates for DSB repair. The resulting “ectopic recombination” between non-allelic TEs can generate chromosomal rearrangements, causing significant fitness reductions in affected individuals (Kupiec and Petes 1988; Symer et al. 2002) and, consequently, the underrepresentation of detected COs around TEs. Longer TEs may present a more significant challenge to the homology required for DSB repair (Waldman 2008) or be more likely to undergo ectopic recombination (Petrov et al. 2003). Yet, our analyses found no associations between the length of TEs without enrichment of H3K9me3 and CO occurrence, suggesting yet other mechanisms mediating the CO suppressive effects of TEs. It is important to note that these potential mechanisms can only arise because we estimated the CO-suppressing impacts of *heterozygous* TEs, which likely resemble the situation in natural populations given the predominantly low population frequencies of TEs (Cridland et al. 2013; Rech et al. 2022).

The observed suppressive effect of TEs on recombination may have a multifold impact on the evolution of both TEs and host genomes. Such an effect could influence TE evolutionary dynamics by limiting their potential for ectopic recombination (Kent et al. 2017; Choi and Lee 2020) and, in turn, shape the evolution of host-mediated epigenetic silencing targeting TEs (Huang and Lee 2024). Given the prevalence of TEs in the gene-rich euchromatic genome, their suppressive effects on recombination could fundamentally influence genic evolution, hindering the fixation of adaptive variants or the removal of deleterious alleles due to selective interference (Hill and Robertson 1966; Felsenstein 1974). Considering the abundance of TEs in *D. melanogaster* (Rech et al. 2022), the estimated 28% TE-mediated reduction in CO occurrence would translate to a roughly 3% decrease in population recombination rate (4Nr or ρ) for the entire euchromatic genome. While this rate may seem modest, the distribution of TEs across the euchromatic genome is not uniform, and some regions may be especially susceptible to such TE-mediated suppressive effects. For example, in genomic regions where TEs preferentially insert (Sultana et al. 2017), ρ may be sufficiently reduced to have a substantial evolutionary impact. In other genomic regions, the influences of TEs on ρ would depend jointly on their suppressive effects on recombination and population frequencies. TEs with moderate enrichment of repressive marks that have minimum fitness impacts (e.g., in intergenic sequences) should have higher population frequencies and thus a greater overall impact on ρ. In either case, the consequentially reduced selection efficacy against TEs due to suppressed recombination should further increase TE population frequencies and drive the accumulation of new TE insertions. This process can further extend and exacerbate the suppression of recombination, eventually resulting in blocks of genomes with diminished recombination rates. These effects may be especially important for generating micro-haplotype that help maintain co-adapted alleles (Connallon and Olito 2022) or the evolution of chromosomes with nearly complete suppression of recombination (Kent et al. 2017; Wei et al. 2020a). It is worth noting that higher TE density (Kim et al. 2021) and stronger TE-mediated spreading of repressive marks (Huang et al. 2022) have been observed in other *Drosophila* species, where the suppressive effects of TEs on population recombination rate could be even more pronounced.

The ability of TEs to copy and move between genome locations make them a significant force shaping the eukaryotic genome. Beyond their previously known role in contributing to varying epigenome (Quadrana et al. 2016; Lee and Karpen 2017) and transcriptome (Cao et al. 2020), our findings reveal that the dynamic mobilome can actively alter the recombination landscapes. This effect contributes to varying recombination within genome, between individuals, and likely across time and even between species, underscoring their critical role in driving genome evolution through diverse mechanisms.

## Materials and Methods

### Drosophila strains used and crosses performed

We chose three DSPR founder strains (King et al. 2012a) that have the maximum differences in SNPs and set up two crosses: A4 x A6 and A4 x A7. These three strains were originally collected from Zimbabwe, Africa (A4), Georgia, USA (A6), and Kenting, Taiwan (A7). To generate F2 recombinants, F1 female offspring from each cross were backcrossed to A4. 192 F2 backcross females were collected for the benchmark experiment, while 7,166 and 7,141 F2 backcross female offspring were collected for A4 x A6 and A4 x A7 crosses, respectively.

### Generation of benchmark data and experimental pool

For the benchmark data, high–molecular weight (HMW) DNA was extracted from 192 females individually using Qiagen MagAttract HMW DNA Kit. The DNA of each individual was divided equally for standard Illumina short–read sequencing and pooled PacBio long-read sequencing. Illumina libraries were generated for individual F2 using Illumina Nextera DNA Flex Library Prep Kit, multiplexed, and sequenced using Illumina NovaSeq 6000 paired-end sequencing (100 bp read length), with at least 5 million reads per library. The sequencing coverage ranges between 4.6X to 54.3X, with an average of 11.5X. The remaining DNA from the 192 individuals were pooled with equal molar ratios for SMRTbell library prep and PacBio CLR sequencing (one SMRT-cell). For the experimental pool, HMW DNA was extracted en masse (∼7000 F2 females) for each cross using Qiagen Blood & Cell Culture DNA Midi Kit. After SMRTbell library prep, libraries were sequenced with PacBio CLR (3 SMRT-cells for A6 library; 2 SMRT-cells for A7 library).

After filtering out reads with low mapping quality (<20) and short length (<4 kb), the mean read length is 22.7kb for the benchmark data, 26.7kb for the A6 cross, and 28.5kb for the A7 cross. The experimental pools have longer PacBio read lengths than the benchmark pool and, therefore, may have higher efficiency in detecting CO events under certain coverages. See **Table S1** for various characteristics of the PacBio sequencing results.

### Annotation of TEs in the assembled genomes of parental strains

For all sequence analysis, we used following genome assemblies from (Chakraborty et al. 2019): NCBI GCA_003401745.1 (A4), GCA_003401885.1 (A6), and GCA_003401915.1 (A7)). To annotate TEs, we ran Repeatmoder2 (version 2.0.3; (Flynn et al. 2020)) with options ‘-engine ncbi -LTRStruct’. The output libraries were used as input for RepeatMasker (version 4.1.0; Smit, AFA, Hubley, R & Green, P. RepeatMasker Open-4.0.2013-2015) to annotate TEs. Repeats classified as LTR, LINE, DNA, or unknown were selected as candidate TEs. To assign candidate TEs to the family level, we blasted TE sequences (using blastn; (Camacho et al. 2009)) to TE sequences annotated in *D. melanogaster* reference genome (version 6.32) and ‘canonical’ TE sequences of *Drosophila* (retrieved from Flybase February 2022) with following parameters: -evalue 1e-5; -perc_identity 80. When a TE was blasted to multiple families, if at least 80% of the query sequence was aligned to one single TE family, the TE was assigned to that family. In all other situations, that TE would be classified as “ambiguous.” TEs located within 5 kb and of the same TE family were merged, while those of different families were excluded from the analysis. We also excluded TEs shorter than 200bp and INE-1, which is mostly fixed in this species (Rech et al. 2022). We used alignment generated by minimap2 (v. 2.24 (Li 2018)) to identify homologous alleles of TE insertions and exclude TEs shared between strains.

### Filtering of regions with residual heterozygosity in the parental strains

Each parental strain (A4, A6, and A7) were resequenced by extracting DNA from 40 female adults using Qiagen DNeasy Blood & Tissue Kit, followed by library prep using Revvity NEXTFLEX Rapid DNA-Seq Kit 2.0 and NovaSeq 6000 sequencing (100bp paired-end sequencing). Illumina short read data were trimmed with Trim Galore (v. 0.6.6) ([CSL STYLE ERROR: reference with no printed form.]) before being mapped to the respective reference genome using bwa mem (v. 0.7.17, (Li 2013)). Mapped reads were sorted using samtools (v. 1.10, (Li et al. 2009)). Following GATK guidelines (v. 4.1.9.0 (McKenna et al. 2010; Van der Auwera et al. 2013)), we employed MarkDuplicates and AddOrReplaceReadGroups, which was followed by using HaplotypeCaller to identify heterozygous sites. Identified within-strain heterozygous sites were removed from all following analyses.

### Identification of COs with short-read sequencing data

We used minimap2 with options -cx asm10 --cs=long, followed by paftools, to call SNPs between two parental genome assemblies. To identify CO in each F2 individual, Illumina short reads were processed with the pipeline described above, with A6 as the reference for mapping. After identifying heterozygous (A6 origin) and homozygous (A4 origin) SNPs in each F2 individual, we used 50-SNP sliding windows to identify the transition in parental origin (i.e., switch between homozygous A4 origin and heterozygous A6 origin). We required the parental tracks to be consistent for at least 200 SNPs to call a CO event. Each identified CO event was visually inspected by plotting the parental tracks along chromosomes.

### Identification of COs from pooled, long-read sequencing data

PacBio CLR reads from each cross were mapped to both parental genomes separately (A4 and A6; A4 and A7) using minimap2. To exclude reads containing potential gene conversion events, we required the aligned regions of a read to be at least 5 kb long and have 20 mapping quality. We used 10-SNP sliding windows along each read to identify transitions in parental origin that are characteristic of CO events. We required that the proportion of SNPs from a particular parent must be at least 50% to assign parental origin. For instance, if the proportion of A4-originated SNPs in a window exceeded 0.5, that window was designated as belonging to the A4 parent. For a read to be considered containing a CO event, each parental haplotype/track needs to cover at least 5 SNPs, and the CO event is at least 2 kb from the edge of the alignment. Reads with more than one switch in parental origin were filtered, due to them likely being caused by sequencing error, gene conversion, or, rarely, double crossovers.

### Assessing potential biases in detecting COs using in silico simulations

To investigate the impact of sequencing error on the accuracy of our approach, we simulated PacBio CLR reads using PBSIM2 (Ono et al. 2021). To assess the false-positive rate, we simulated 100x read depths from each of the A4 and A6 genomes without recombination, with parameter --data-type CLR --model_qc model_qc_clr --length-mean 18000 --length-sd 2500, a condition similar to that of the benchmark data. Then, we applied our method to identify any false-positive CO events from these reads. To assess the impacts of sequencing errors on the sensitivity of our algorithm, we simulated PacBio CLR reads from 7,000 simulated recombinants (between A4 and A6), mimicking the real pool. In these simulations, one CO was randomly placed in each chromosome arm (five CO events per genome). The average coverage across the recombinant genomes was set to be 0.1 to mimic the A6 experimental pool. For identifying simulated reads covering COs, we employed the same criteria as those used in the experimental pool analysis. The proportion of reads covering CO but were not identified by our criteria is the *false-negative rate per read*.

To explore how sequencing depth impacts the recall rate of COs, we simulated a pool of 400 recombinants with one CO per chromosome arm (2,000 CO events in total) and 5,000X PacBio CLR reads from that pool. After reads alignment, we downsampled reads to average 8X, 6X, 4X, 2X, 0.5X, 0.25X, and 0.1X per recombinant genome. We then used the same criteria to identify CO events and estimated the CO recall rate for different read depth.

### CUT&Tag experiment and analysis

To profile the distributions of H3K9me3 in the A6 and A7 parental genomes, we collected 16-18hrs-old embryos laid by 2-7-day-old adults after one hour of pre-laying. Nuclei were prepared from fresh embryos, and ∼100,000 nuclei were used per biological replicates, with three biological replicates per strain. We followed the CUT&Tag@home protocol (V.3, (Kaya-Okur et al. 2019; Henikoff et al. 2020)) using H3K9me3 rabbit antibody (Abcam EPR16601) and anti-rabbit Secondary Antibody binding (EpiCypher 13-0047). After the pA-Tn5 adapter complex binding and library prep steps, the libraries were analyzed by Tapestation and followed by Illumina sequencing (Illumina NovaSeq 6000, 100 bp paired-end sequencing). Due to the challenges of performing epigenomic analysis with meiotic cells, CUT&Tag was performed using developing embryos. The epigenetic silencing of TEs is directed by piRNA (Sienski et al. 2012; Le Thomas et al. 2013), whose profiles are highly correlated between ovaries (containing meiotic cells) and embryos (Brennecke et al. 2008). Furthermore, the generation of piRNAs in embryos depends on those inherited maternally (Brennecke et al. 2008; Le Thomas et al. 2014), both suggesting that H3K9me3 enrichment around TEs observed in embryos is a good proxy for those in meiotic cells.

The generated data were analyzed following CUT&Tag Data Processing and Analysis Tutorial (Henikoff et al. 2020; Zheng et al. 2020). In short, Illumina short reads were trimmed using Trim Galore (v. 0.6.6), then mapped to A6 or A7 genomes respectively with bowtie2 (version 2.4.5, (Langmead et al. 2009)), and filtered for at least 20 mapping quality using samtools (v. 1.10, (Li et al. 2009)). The histone-modification magnitude (HM) for each position was estimated as the sequencing coverage using bedtools (v2.30.0, (Quinlan and Hall 2010)). Only read pairs on the same chromosome with fragment lengths smaller than 1,000bp were kept for the following analyses.

We used two measures to quantify H3K9me3 enrichment around each TE: adjacent H3K9me3 enrichment and H3K9me3 total mass. For adjacent H3K9me3 enrichment, we calculated the mean HM for 5 kb upstream and downstream flanking each TE. This estimate was then normalized by the mean HM of 20–40 kb upstream and downstream of the respective TE, following (Lee and Karpen 2017). Normalized HM above one is deemed enriched with H3K9me3. The H3K9me3 total mass is the accumulated K9 enrichment level across the extent of H3K9me3 spreading from a TE. The extent of H3K9me3 spreading was inferred by first dividing the 20 kb upstream and downstream from a TE into 1 kb nonoverlapping windows, estimating normalized HM for each 1 kb window, and identifying the farthest window from TEs in which the normalized HM was consecutively above one. The total K9 mass is the sum of normalized HM across 1 kb windows within the extent of the spreading.

### Indexes used to investigate TEs’ impacts on CO occurrence

We used two indexes: the number of COs around a TE insertion (CO number) and the distance from a TE to the nearest CO (distance to CO). For the CO numbers, we required both 5 kb windows to contain at least five diagnostic SNPs, and both immediately neighboring SNPs of CO events needed to be located within the window. For control windows, we selected two adjacent 5 kb windows (10 kb) that are at least 30kb away from any TEs and have at least 5 SNPs and assigned CO events to either window with the same criteria. For distance to CO, we identified the nearest CO upstream/downstream to a focal site (TE or control site) and used the shorter of the two.

### Testing the associations between the presence of TEs and CO occurrence *within* strains

We used one-tailed *Mann-Whitney U tests* to compare the CO numbers between TE and control windows with TEs as well as glm models to analyze the CO numbers with the Negative binomial family function:

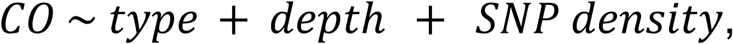

where *type* is whether a window is with or without TE, while *depth* and *SNP density* are sequencing depth and number of SNPs in the flanking windows. To compare the effects between TEs with and without H3K9me3 enrichment, we used H3K9me3 enrichment level in the windows and performed glm (*CO ∼ H3K9me3enrichment*) and *Spearman rank correlation tests*.

For the distance to the nearest CO (*D*), we also performed *Mann-Whitney U tests* and linear regressions:

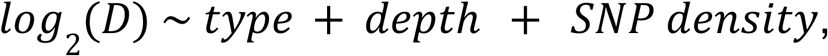

where *type* is whether a window is with or without TE, while *depth* and *SNP density* are sequencing depth and number of SNPs in the region between the focal site and the nearest CO. To compare the effects between TEs with and without H3K9me3 enrichment, we used H3K9me3 mass and performed linear regression (*CO ∼ H3K9me3 mass*) and *Spearman rank correlation tests*.

### Testing the associations between the presence of TEs and CO occurrence *between* strains

For analysis comparing CO numbers, we selected control windows based on the following criteria: (1) at least 10kb away from TEs in either strains, (2) neither homologous alleles show enrichment of H3K9me3 (< 1), and (3) the ratio of SNP density between homolog windows is similar to that of TEs. For (3), we binned control window pairs using ten quantiles of the SNP density ratio between homolog windows of TEs and then downsampled bins with excessive window pairs to have the same number of control windows across bins. For TE and control windows, we downsampled reads to have matching read depth between homologous alleles to control for the effect of sequencing depth on the detection of COs. We used two approaches to test whether the differences in COs (with TE allele minus without TE allele) differ between TE and control windows. The first approach downsamples sequencing depth for TEs and control windows, performs one-tailed *Mann-Whitney U test,* and repeats the process 1,000 times to get a median *p* for the *Mann-Whitney U test.* The second approach bootstraps control windows to the same numbers as TE windows per arm, downsamples a bootstrapped set, estimates the mean difference in CO number for each downsampled sets, and generates a null distribution of the median of the mean differences in CO number for 1,000 bootstrapped sets. One-tailed *p-values* were then estimated by comparing the median of mean differences in CO for downsampled TE windows.

For analysis based on the distance to the nearest CO, we also employed downsampling to ensure equal sequencing coverage between the homologous alleles. During the process, if the nearest CO was unsampled, the next nearest CO was used to estimate the distance to the nearest CO. All other analysis procedures are the same as those for analysis using CO numbers.

To test the impact of TE-induced H3K9me3 on the occurrence of COs, we classified TEs into those with and without H3K9me3 enrichment (adjacent H3K9me3 enrichment > 1 or not) for analysis using CO number or high/low H3K9me3 mass (use 50% quantile of H3K9me3 mass) for analysis using distance to nearest CO. TEs whose non-TE homologous alleles have adjacent H3K9me3 enrichment > 1 (for analysis using CO number) or H3K9me3 mass > 3 were excluded. We used the same two approaches as above to investigate whether TEs with H3K9me3 enrichment show significantly stronger suppression on CO occurrence, with the only difference being the null distribution is now generated from bootstrapped non-H3K9me3 enriched TEs. For *Spearman correlation tests* between H3K9me3 enrichment and differences in CO numbers/ratio of distance to nearest COs, we used the median correlation coefficient and *p-values* of 1,000 downsampled sets.

### Using DSPR RILs to test the effect of TEs on CO occurrence

The DSPR RILs belong to a multiparental mapping panel that was created from a 50-generation intercross between eight inbred founder lines, followed by 25 generations of inbreeding (King et al. 2012a, 2012b). In other words, the panel of RILs consists of inbred lines whose genomes are mosaics of the original eight parental lines. Previously, King et al. (King et al. 2012a, 2012b) developed a hidden Markov model to infer the probability of founder ancestry at each genomic location within each DSPR RIL. This model was implemented using genome sequencing data for the founder lines and genotyping data at ∼10,000 positions for the full set of DSPR RILs, resulting in highly accurate founder assignments across most of the genomes of the full panel of RILs.

A subset of the DSPR RILs (28 RILs) have whole-genome Illumina sequencing data available (NCBI SRA PRJNA1161589; (King et al. 2012b)). These data were aligned to the *D. melanogaster* release 5 reference genome with bwa mem, and SNPs were called using samtools mpileup. We used these data with the previously developed hidden Markov model (King et al. 2012a) to infer the underlying founder ancestry at each genomic position in each RIL with high precision and accuracy. For each RIL, we defined every breakpoint interval where the founder assignment transitions from being 95% probable for one founder line to 95% probable for a different founder line, thus inferring a CO event at this interval. COs present in multiple RILs (i.e., with the same parental strains generating the COs and the same CO locations) were filtered to avoid double counting. We also removed COs in heterochromatin. At the end, 1,801 euchromatic COs were used for the analysis.

To identify TEs in DSPR founders, we used SV identified in (Chakraborty et al. 2019) and the above approach to identify and annotate TEs into families. The same TE insertion site with more than one TE family called in different strains was filtered. We also removed INE-1 TEs. These filtering criteria yielded 4,686 euchromatic TE insertions. To be included in the following analysis, we further required a TE insertion site to have at least two with TE and two without TE alleles. TE alleles are those where TEs are present in one of the two strains from which the CO was generated. Accordingly, an allele may be considered as a TE allele even if that particular TE is not present in the RIL. All other alleles were considered without TE alleles. We then estimated the median physical distance (in bp) between a TE insertion site and the nearest COs for alleles with and without TEs.

### Generation of transgenic strains with candidate TEs and mCherry marker

pAct5C promoter and the mCherry coding sequences were synthesized and cloned into pBS-KS-attB1_2 vector (obtained from DGRC, stock number 1322; (Venken et al. 2011)) by Genscript (Piscataway, NJ). Plasmid with TE sequences downstream of mCherry was obtained by cloning PCR amplified transposons into BamH1-digested pBS-KS-attB1-2 _pAct5C_mCherry plasmid using Takara infusion technology (https://www.takarabio.com). 1360 was amplified from genomic DNA into one fragment, and Roo and Copia were amplified into two fragments from BAC clones obtained through Drosophila Genomic Resource Center (DGRC) (CH321-11A21 for Roo and CH321-23O24 for Copia) with the following primers.

Primers used for amplifying TEs For 1360:

- Forward: tcttatcgagggatcgaaaggaatacggtattaccaaag
- Reverse: cgacaagcttggatccatcggttgatgatcaataaatttc

For Roo:

- Fragment 1 Forward: tcttatcgagggatcatcgatatattcgtgttcatgtgtgaacattc
- Fragment 1 Reverse: gattccacgattaccttagcatggat
- Fragment 2 Forward: ggtaatcgtggaatcactccaagca
- Fragment 2 Reverse: cgacaagcttggatcatcgattgttcacacatgaacacgaatatatt

For Copia:

Due to numerous repeats in the Copia causing non-specific primer binding, nested PCR was performed. To overcome this problem, PCR fragments were initially obtained using primers outside of the Copia, followed by nested PCR on the obtained fragments. The primers used for Copia amplification are:

For the initial PCR:

- Phage Forward Primer: gtcagtcgatggggaaatgcag
- Fragment 1 Reverse: cattataggatatctgaggcttagtctttaatctct
- Fragment 2 Forward: agatatcctataatgaagaggataatagtctaaataaagttgttctaaatg
- Phage Reverse Primer: tcagcagcaaaacgaagcatgt

For nested PCR:

- Fragment 1 Forward: tcttatcgagggatcatcgattgttggaatatactattcaacctacaaaaataacgttaaa
- Fragment 1 Reverse: cattataggatatctgaggcttagtctttaatctct
- Fragment 2 Forward: agatatcctataatgaagaggataatagtctaaataaagttgttctaaatg
- Fragment 2 Reverse: cgacaagcttggatcatcgattgttgtttaacgttatttttgtaggttgaa

All plasmids (mCherry only, mCherry + 1360, mCherry + Roo, and mCherry + copia) were confirmed by sequencing and injected into MIMIC strain with docking site at 2L:17,325,574 (y[1] w[*]; Mi{y[+mDint2]=MIC}MI02983; BDSC 48358) by BestGene (Chino Hills, CA).

### Estimation of recombination rates around transgenic TE insertion

To estimate recombination rate around the construct insertion sites, we used the approach developed in (Wei et al. 2020b), which infers recombination rate based on shifts in allele frequencies along the chromosome. We crossed each transgenic strain to *b/b* homozygotes (BDSC 227) and collected ∼1,000 mixed-sex F2 backcrossed offspring with *b/b* genotype. Backcrossed *b/b* offspring (mixed sexes) were pooled with ∼1000 individuals for each genotype. DNA was extracted from the selected F2 as a pool with a Qiagen DNeasy Blood & Tissue Kit, following the protocol in (Wei et al. 2020b). The pooled DNA was sheared by Covaris S220, targeting 300-400 bp, prepped as sequencing libraries with Revvity NEXTFLEX Rapid DNA-Seq Kit 2.0, and sequenced with Illumina NovaSeq 6000 (150 bp paired-end sequencing). The median sequencing depth is ∼60-88 across pools. The experiment was conducted in three blocks, with the first block containing mCherry, mCherry+1360, and mCherry+Roo, while the rest of the blocks contained all genotypes. We also sequenced the two parental strains (BDSC 227 and BDSC 48358) by extracting DNA from 40 female adults using Qiagen DNeasy Blood & Tissue Kit, followed by library prep using Revvity NEXTFLEX Rapid DNA-Seq Kit 2.0 and Illumina NovaSeq 6000 sequencing (150 bp paired-end sequencing).

Sequencing data were then analyzed using the analytical pipeline described in (Wei et al. 2020b). In short, sequences of the pools and parents were aligned using bwa mem to the reference *D. melanogaster* genome release 6 and genotyped using GATK’s recommended pipeline. To procure a set of high confidence diagnostic sites, we removed sites with GQ < 30 in the parental genotypes and isolated positions where the parents have different homozygous alleles. We then removed sites with coverage beyond 5th and 95th percentile. The final diagnostic sites were then used to phase alleles in the marker-selected pools. Allele frequency at every site was then infrared from the Allele Depth field in the vcf. Allele frequency across the chromosome was smoothed using local regressions with LOESS. After removing the paternal contribution, recombinant fractions were then estimated.

## Data availability

All the sequence data generated in this study was deposited to SRA under the accession number PRJNA1159377. The scripts for performing the analyses are available at https://github.com/YuhengHuang87/TE_crossover and https://github.com/weikevinhc/Marsupial

## Supporting information

STables

SFigures

## Acknowledgment

We appreciate Charles Langley for suggesting the use of PacBio sequencing, Serafin Colmenares for suggestions on the design of transgenic constructs, Stephen Wright, Andrea Betancourt, Tony Long, J. J. Emerson, and Harsh Shukla for helpful discussion of the project, and Ching-Ho Chang for critically reading the manuscript. We thank the University of California High-Throughput Genomics Facility and High-Performance Cluster at UC Irvine for sequencing and computational resources. This work was supported by F32 GM099382 to EGK, the European Research Council (ERC) under the European Union’s Horizon 2020 research and innovation program, grant agreement No. 850405 and VIDI VI.Vidi.203.001 financed by the Dutch Research Council (NWO) to AJ, NSERC Discovery Grant (RGPIN-2023-05390) to KHCW, and NIH R35GM142494 to YCGL.

