## Supplementary material for "Varying recombination landscapes between individuals are driven by polymorphic transposable elements": SFigures

**Figure S1. Resolution of CO breakpoints for benchmark data and real experimental pool.** CO events were narrowed down to between two SNPs, where the parental origin of haplotypes switched. The resolution is determined by SNP density, sequencing error, and the molecule produced by the recombination event. Most CO events could be narrowed down to within 1 kb. There is no significant difference in resolution between benchmark data (A) and either of the experimental pools (B). *Mann-Whitney U test*, $p > 0.4$ for all three comparisons.

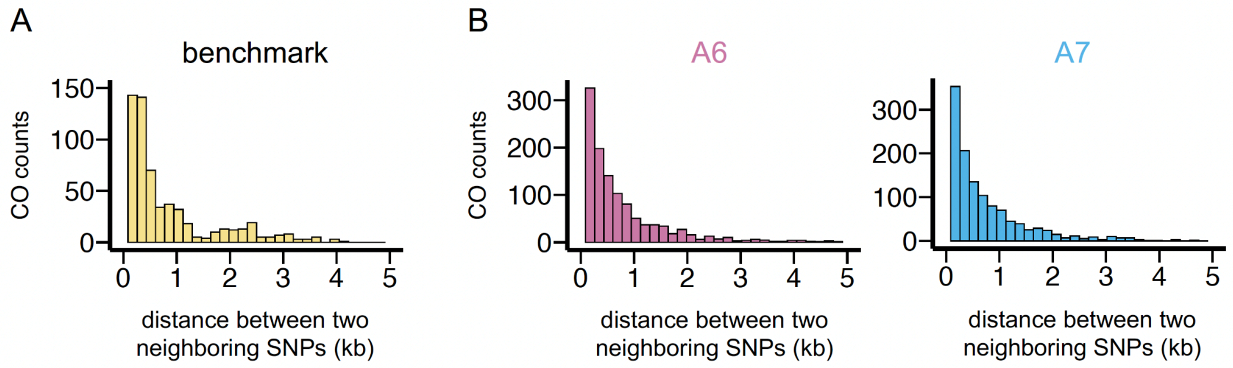

**Figure S2. Distribution of COs identified and missed by the pool-sequencing approach in benchmark data.** To examine whether COs that are missed by the pool-sequencing approach distribute nonrandomly across the genome, we binned nearby CO events identified by the short-read approach into bins 10, 15, or 20 CO events (corresponding to 56, 35 or 28 bins) across the genome. We then performed Fisher's Exact Test to investigate whether any bins have higher false negative rates than the rest of the chromosome arm. No bins have significant enrichment of false negative CO events (FDR adjusted $p > 0.09$), suggesting the random distribution of false negative events.

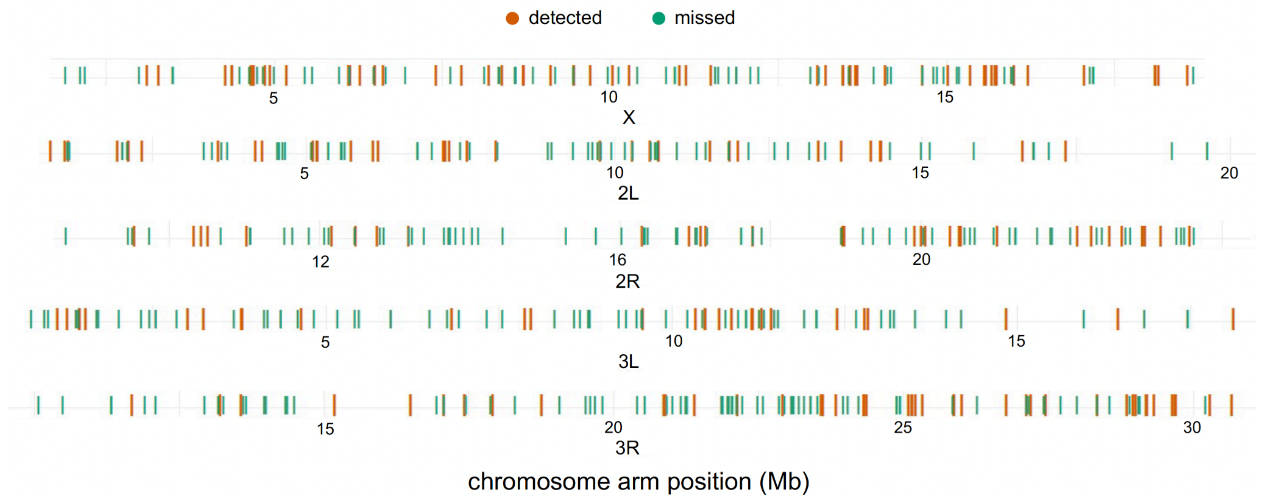

**Figure S3. Distribution of normalized CO numbers across the euchromatic genome in the experimental pools and previously reported recombination map (Comeron et al. 2012).** A sliding window of 1 Mb with a step size of 100 kb was used to plot the Comeron et al. 2012 recombination map (top) and the depth-normalized CO numbers in this study (bottom).

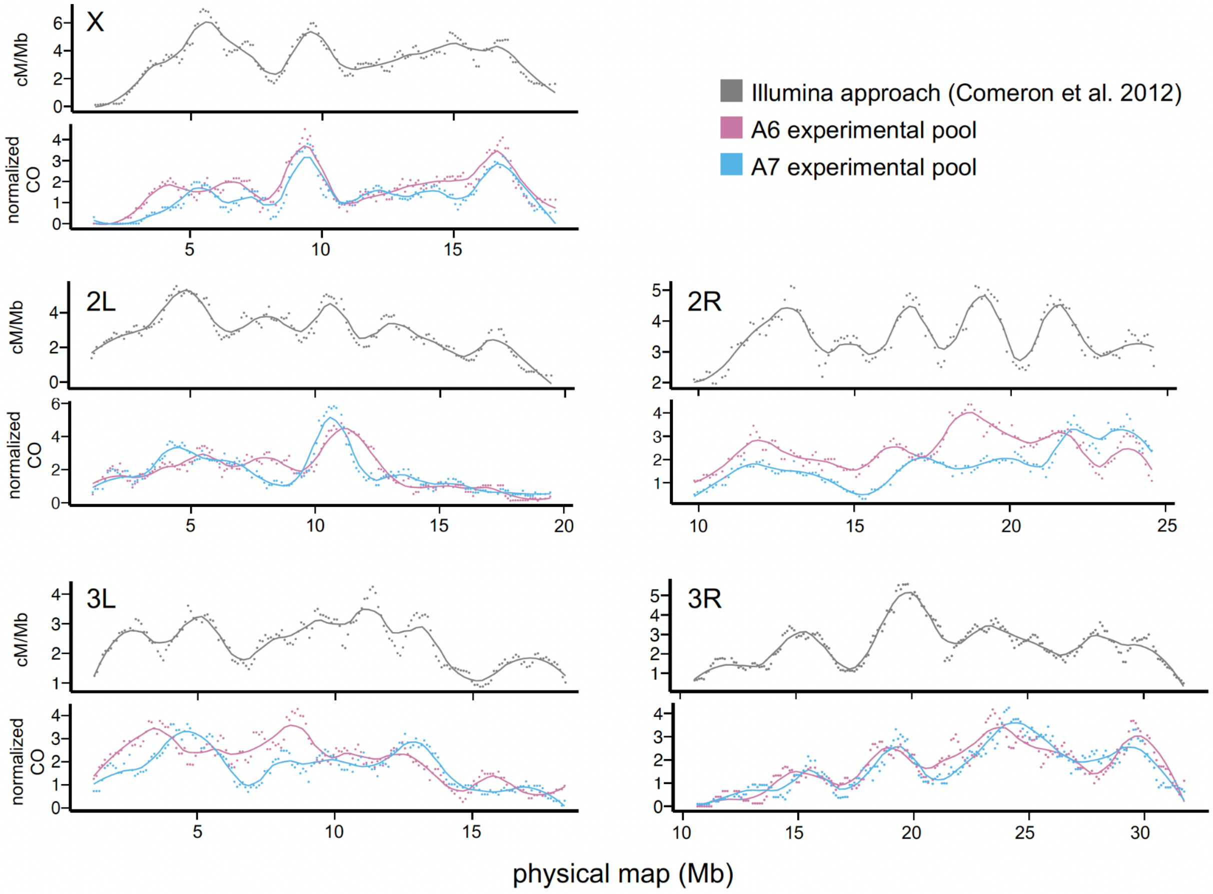

**Figure S4. Distribution of CO events that were identified and missed in simulations with respect to SNP density.** By binning 20 nearby CO events to estimate the local false negative rate, we found one euchromatic region with a significantly higher false negative rate than the rest of the chromosome arm (X:1272264..2062069, FDR adjusted $p = 0.004$). This region also has low SNP density, suggesting the lack of SNPs may explain the high false negative rate. The SNP density was plotted as the number of SNPs between A4 and A6 genomes per 100 kb.

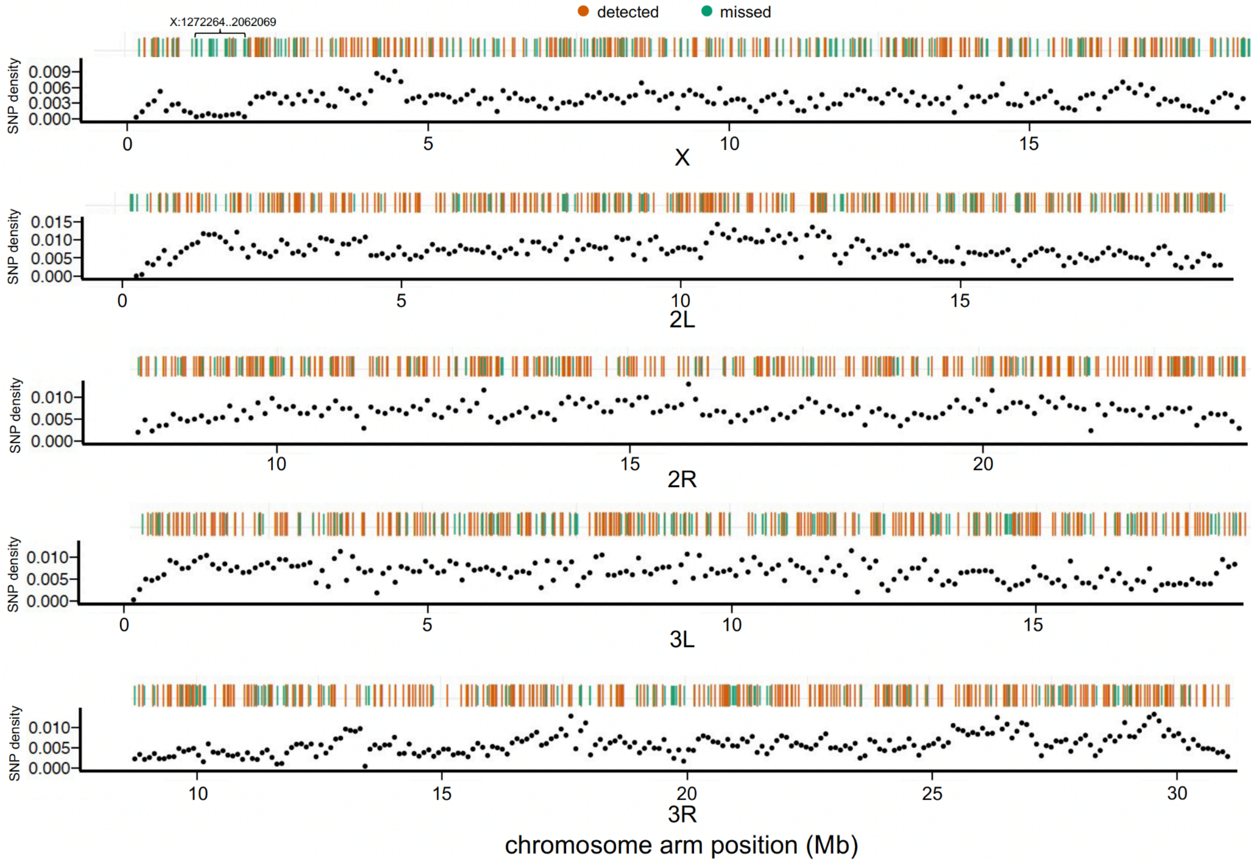

**Figure S5. Assessing the impact of repeated and unequal sampling of COs in the benchmark data and experimental pool.** (A) The curved line represents the expected proportion of repeatedly sampled haplotypes, calculated using multinomial sampling. This calculation assumes an equal DNA contribution from each individual across various depths per haplotype. With the observed depth-to-haplotype ratio of the benchmark data (orange line), the majority (68.4%) of the COs are expected to be repeatedly sampled. In contrast, only 5.2% (A6 experimental pool) and 6.5% (A7 experimental pool) haplotypes are expected to be repeatedly sampled due to the much lower individual-to-depth ratio. (B, C) The distributions of the number of reads supporting a CO for the benchmark data and experimental pool. (B) We compared the distributions of the number of reads supporting each CO event for observed (yellow) and simulated (gray) data for the benchmark pool. The simulated data assume equal pooling of DNA of F2 recombinant, and each CO has the same probability of being sampled. The observed distribution of reads per CO is significantly more right-skewed than expected, suggesting that the F2 DNA was likely pooled unequally. (C) In contrast, few COs were sampled by multiple reads in the experimental pools, suggesting a minimum impact of unequal sampling of F2 DNA. It is worth noting that multiple reads supporting the same CO event can result from repeated sampling of the same CO event or from different CO events located within the same two neighboring SNPs.

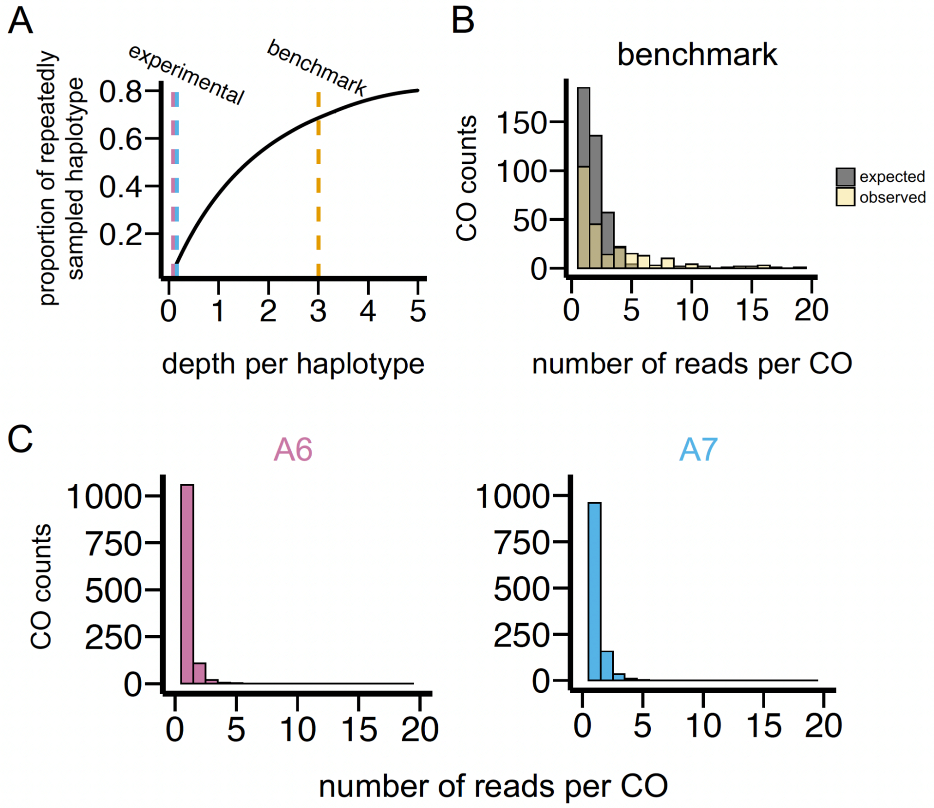

**Figure S6. Sequencing depth and SNP number for TE and control windows in experimental pools in the within-strain analysis).** The sequencing depths for TE windows are significantly lower than control windows in both strains. The SNP numbers for TE windows are significantly lower than control windows in the A6 experimental pool. *Mann-Whitney U test,* *$p < 0.05$, ***$p < 0.001$.

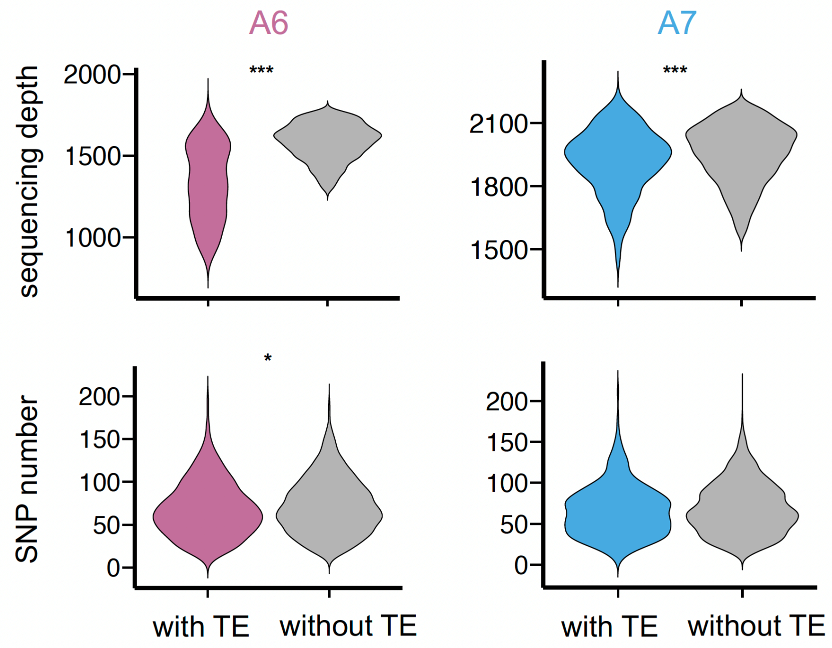

**Figure S7. Quantifications of the enrichment of H3K9me3 around euchromatic TEs.** (A) Average H3K9me3 enrichment level at homologous sequences with and without the presence of TEs. The enrichment is normalized to the local regions 20-40 kb from TEs. Mean H3K9me3 was estimated for every 1 kb window from TEs and smoothed with LOESS (span = 20%). (B) Cartoon illustrating the estimations of the adjacent H3K9me3 enrichment and H3K9me3 mass. The adjacent H3K9me3 enrichment is the mean H3K9me3 enrichment in the 5 kb windows immediately adjacent to TEs (average over both sides). The H3K9me3 mass is the accumulated H3K9me3 enrichment level across the extent of its spreading from a TE. We estimated the extent of H3K9me3 enrichment based on whether the enrichment was consecutively above one for the 1 kb windows starting from TE boundaries (up to 20 kb), followed by summing the H3K9m3 enrichment across the 1 kb windows.

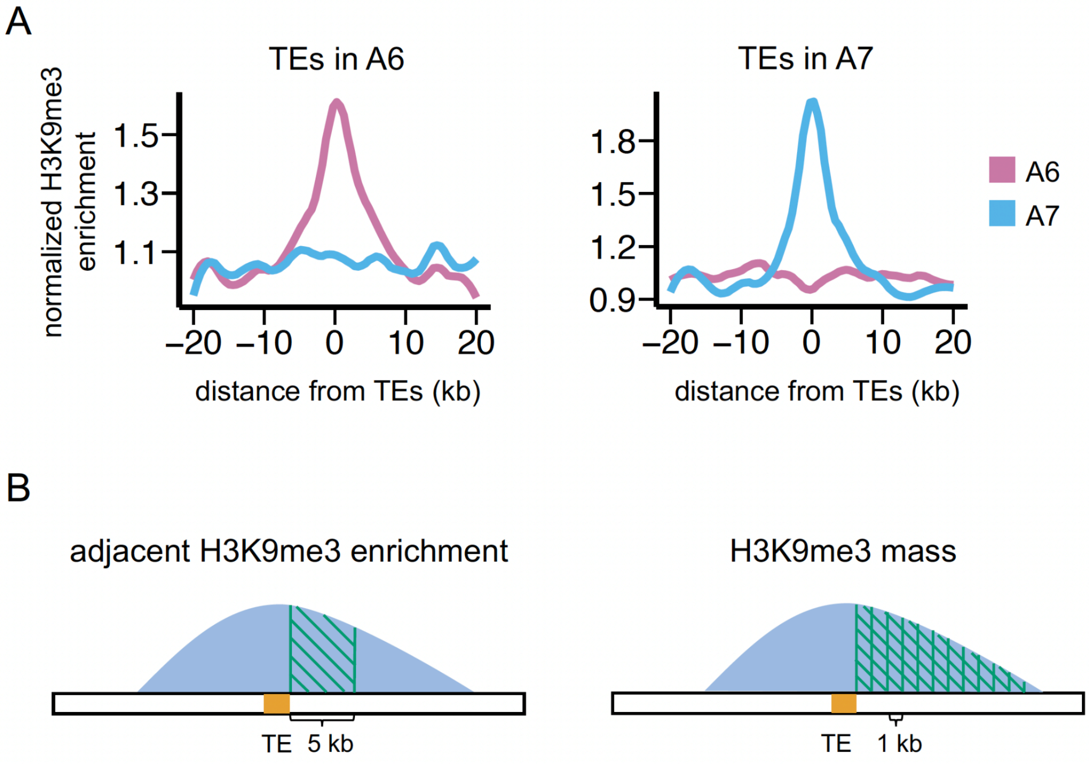

**Figure S8. Impact of TEs on CO occurrence inferred by the between-strain analysis.** (A) The distributions of the *p-values* of the *Mann-Whitney U test* comparing two indexes (difference in CO number and difference in distance to nearest CO) between windows with and without TEs across 1,000 downsampled sets. (B) The distributions of the mean differences in CO number and mean ratio of distance to nearest COs generated by bootstrapping windows without TEs. Dash lines indicated observed values for windows with TEs.

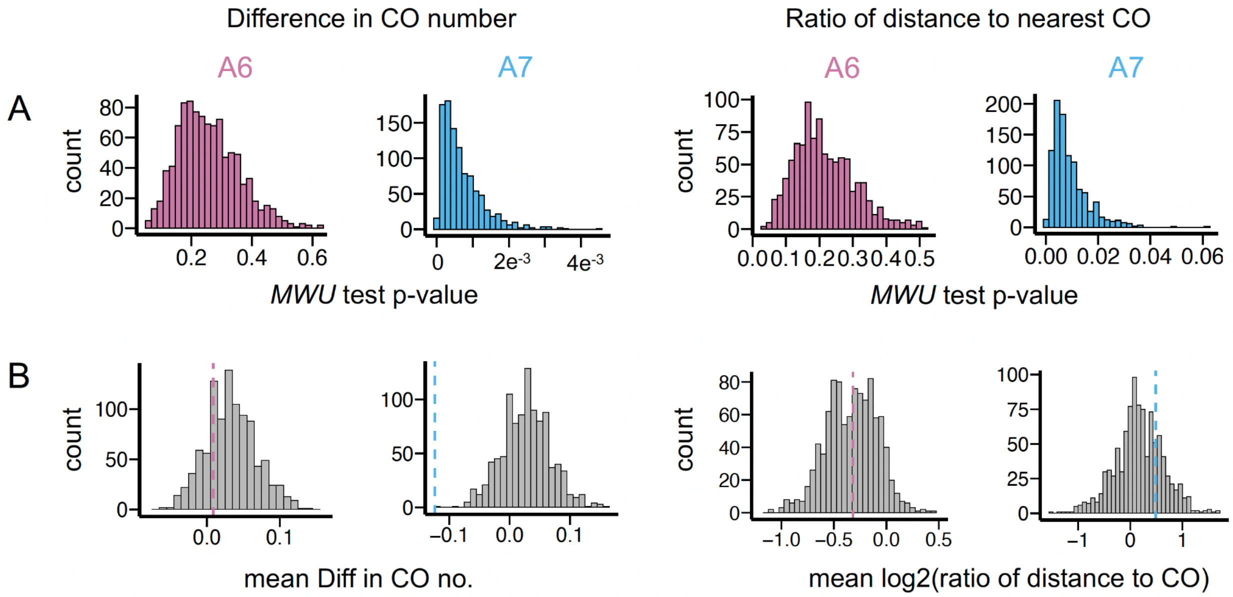

**Figure S9. Impact of TEs with H3K9me3 enrichment on CO occurrence inferred by the between-strain analysis.** (A) The distributions of the *p-values* of *Mann-Whitney U test* of comparing two indexes (difference in CO number and difference in distance to nearest CO) between TE windows with or without H3K9me3 enrichment across 1000 downsampled sets. (B) The distributions of the mean differences in CO number and mean ratio of distance to nearest COs generated generated by bootstrapping the TE windows without H3K9me3 enrichment. Dash lines indicated the estimates from the windows with TEs.

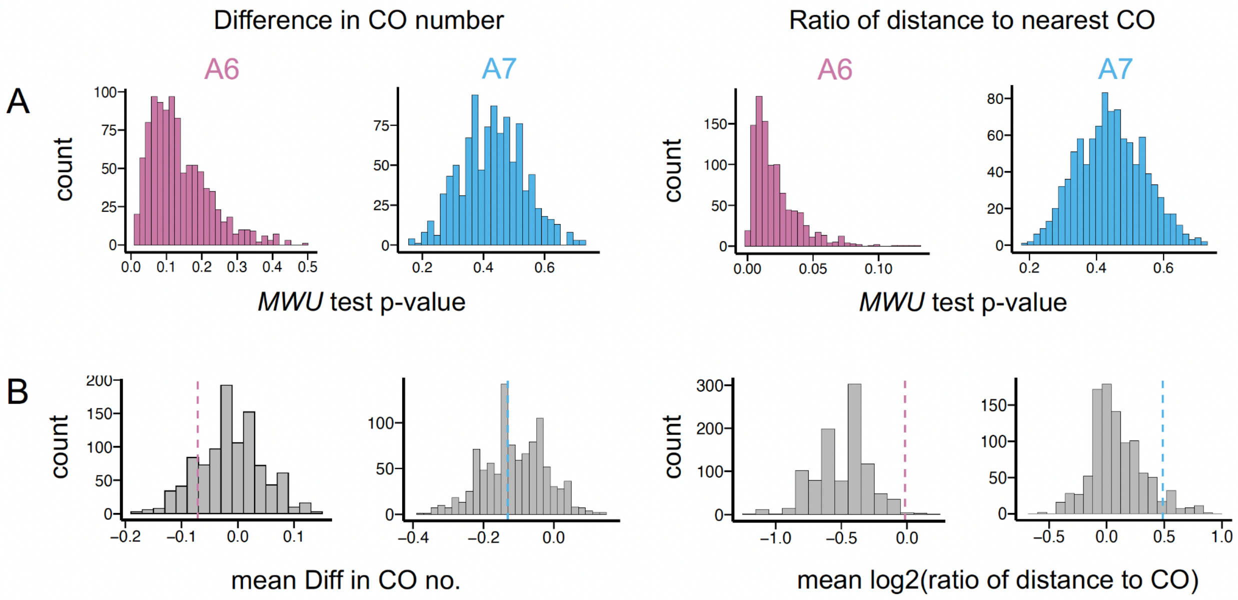

**Figure S10. Correlations between CO occurrence and TE-mediated H3K9me3 enrichment inferred by between-strain analysis.** The distributions of *Spearman correlation coefficients* between the adjacent H3K9me3 enrichment and difference in CO number (left) and between H3K9me3 mass and ratio of the distance to nearest CO (right) for 1000 downsampled datasets.

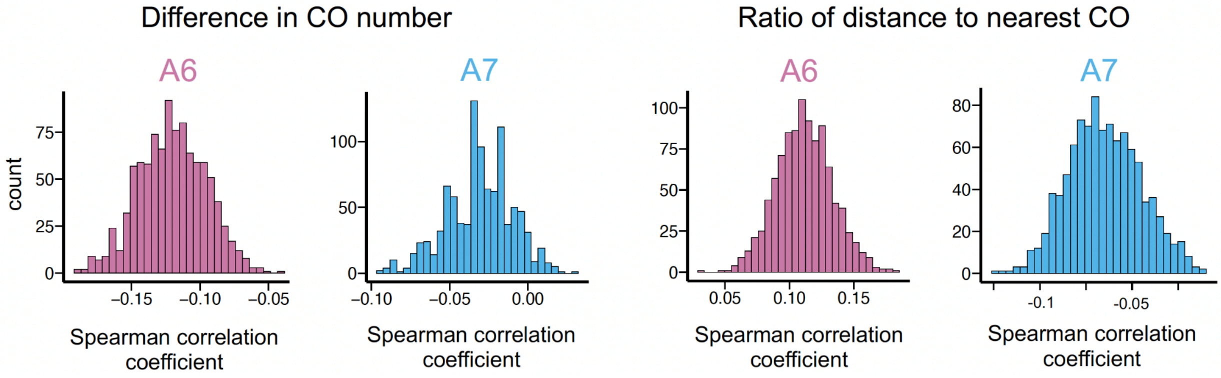

**Figure S11. Correlations between CO occurrence and TE length inferred by between-strain analysis.** The distributions of *Spearman correlation coefficients* between TE length and difference in CO number (left) and between TE length and the ratio of distance to nearest CO (right) for 1000 downsampled datasets.

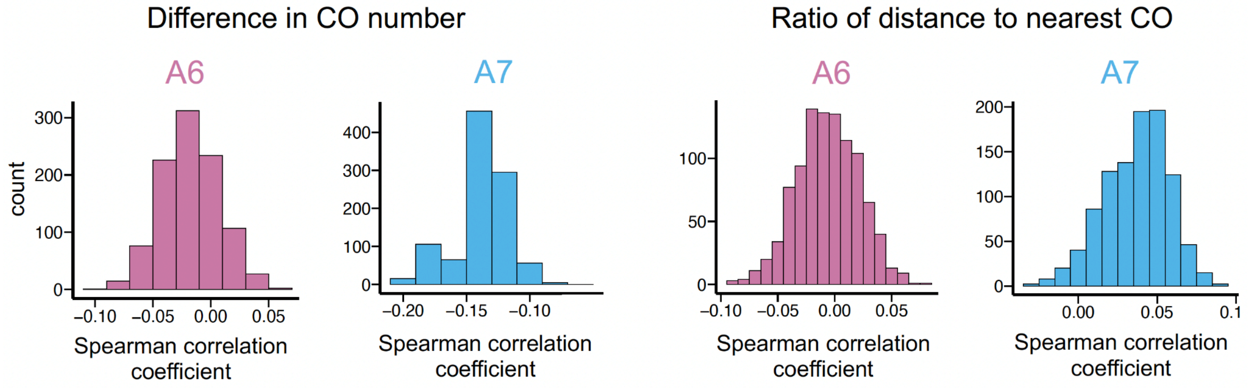
